# DGTS overproduced in seed plants is excluded from plastid membranes and promotes endomembrane expansion

**DOI:** 10.1101/2024.07.11.603045

**Authors:** Sarah Salomon, Marion Schilling, Catherine Albrieux, Grégory Si Larbi, Pierre-Henri Jouneau, Sylvaine Roy, Denis Falconet, Morgane Michaud, Juliette Jouhet

## Abstract

Plants and algae must adapt to environmental changes, facing various stresses that negatively impact their growth and development. One common stress is phosphate (Pi) deficiency, which is often in limiting quantity in the environment. In response to Pi deficiency, these organisms increase Pi uptake and remobilize intracellular Pi. Phospholipids are degraded to provide Pi and replaced by non-phosphorous lipids, such as glycolipids or betaine lipids. During the evolution, seed plants lost their capacity to synthesize betaine lipid. By expressing BTA1 genes, involved in the synthesis of diacylglyceryl-N,N,N-trimethyl-homoserine (DGTS), from different species, our work shows that DGTS can be produced in seed plants. In Arabidopsis, expressing BTA1 under a phosphate starvation-inducible promoter resulted in limited DGTS production without having any impact on plant growth or lipid remodeling. In transient expression systems in *Nicotiana benthamiana*, leaves were able to accumulate DGTS up to 20 % of their glycerolipid content at a slight expense of galactolipid and phospholipid production. At the subcellular level, we showed that DGTS is absent from plastid and seems to be enriched in endomembrane, driving an ER membrane proliferation. Finally, DGTS synthesis pathway seems to compete with PC synthesis via the Kennedy pathway but does not seem to be derived from PC diacylglycerol backbone and therefore does not interfere with the eukaryotic pathway involved in galactolipid synthesis.

## Introduction

Plants and algae are organisms that must constantly adapt to changes in their environment, and thus cope with numerous stresses such as nutrient, water and thermal stress, or attack by various types of pathogen. These stresses can have negative consequences on the growth and development of these organisms, and can therefore have a considerable impact on the yield of raw material production. Phosphate (Pi) deficiency is one of the most common nutrient stresses, leading to a significant reduction in yield - ranging from 25% to 60% in various crops (Nelson and Janke, 2007; Smit et al., 2009; Vitousek et al., 2009). Pi is an essential nutrient for the survival of all living organisms. Plants and algae draw the Pi they need for their development from the soil or aquatic environment. However, although abundant, this nutrient is not evenly distributed and is often present in forms that cannot be assimilated by plants (Tiessen, 2008). This low Pi availability is often compensated by the use of phosphorus-enriched fertilizers, which are subsequently responsible for significant soil and water pollution. In order to limit the use of this type of fertilizer while improving crop yields, it is essential to understand the mechanisms involved in the adaptation of photosynthetic organisms to Pi deficiency (Cuyas et al., 2023).

In the event of Pi deficiency, photosynthetic organisms will set up adaptation mechanisms to increase Pi uptake from the environment and remobilize intracellular Pi reserves. In cells, Pi is mainly stored in the vacuole, but is also present in significant quantities in various biomolecules such as nucleic acids and certain lipids (Poirier and Bucher, 2002). A third of intracellular Pi is present in plant phospholipids, the main constituents of extra- plastidial cell membranes (Poirier et al., 1991). In the event of Pi deficiency, a part of the membrane phospholipids is degraded to release Pi that are then replaced by non-phosphorous lipids to maintain the integrity of cell membranes (Hölzl and Dörmann, 2019; Jouhet et al., 2004; Van Mooy et al., 2009). When phosphate is limiting in plants, glycolipids can replace phospholipids in cell membranes: sulfoquinovosyldiacylglycerol (SQDG) replaces phosphatidylglycerol (PG) in photosynthetic membranes, and digalactosyldiacylglycerol (DGDG) replaces phospholipids in extraplastidial membranes (Andersson et al., 2005; Bolik et al., 2022; Jouhet et al., 2004). However, even after prolonged Pi deficiency, only a fraction of the phospholipids is degraded, limiting intracellular Pi remobilization and halting cell growth, probably because the physicochemical properties of phospholipids and glycolipids are different (Kanduč et al., 2017).

Algae, as well as some fungi, bacteria or bryophytes, are able to synthesize another type of non-phosphorous glycerolipid called betaine lipid. Currently, three species of betaine lipids have been described in photosynthetic organisms: diacylglyceryl-N,N,N-trimethyl-homoserine (DGTS), diacylglycerohydroxymethyl- N,N,N-trimethyl-b-alanine (DGTA), and diacylglycerylcarboxy-N-hydroxymethylcholine (DGCC) (Dembitsky, 1996; Kato et al., 1996; Sato, 1992). In algae, it is often suggested in the literature that these two lipid classes (i.e. phospholipids and betaine lipids) are interchangeable, because like phospholipids, betaine lipids are synthesized and localized in extraplastidial membranes (Eichenberger et al., 1993; Künzler et al., 1997), and because they share a trimethyl-nitrogen fragment with phosphatidylcholine (PC) (Eichenberger et al., 1993; Kato et al., 1996). Indeed, *Chlamydomonas reinhardtii* has no PC but only DGTS (Yang et al., 2004), and in the yeast *Saccharomyces cerevisiae*, PC can be replaced by DGTS without major impact on cell growth (Riekhof et al., 2014). However, looking across a large number of organisms, no correlation could be found between DGTS and PC content (Künzler and Eichenberger, 1997). The complementary association between betaine lipids and phospholipids has long been accepted as an adaptive strategy to cope with periodic phosphate deficiency in the natural environment. However, the degree of change in cellular betaine lipids and PC contents upon fluctuations of the external phosphate (Pi) availability is quite different among species (Cañavate et al., 2017) and finally betaine lipid physicochemical properties are not so similar to PC as inferred (Bolik et al., 2023).

These species-dependent differences imply metabolic diversity among algae, as well as versatility in betaine lipids function and requirement under Pi-starved conditions (Cañavate et al., 2016), yet we lack a comprehensive understanding of betaine lipids functions. Despite this need for extending research on betaine lipids from microalgae, only a few contributions followed the initial works compiled by (Dembitsky, 1996). The presence and the description of betaine lipids in cells are only based on biochemical analyses and therefore poorly cover the wide range of organisms (Cañavate et al., 2016; Dembitsky, 1996; Künzler and Eichenberger, 1997; Makewicz et al., 1997; Vaskovsky et al., 1998). In a simplified view, betaine lipids are present in lower plants, algae and some fungi but even these groups show several exceptions and they are absent from seed plants, i.e. gymnosperm and angiosperm (Rozentsvet, 2004). All these studies were accomplished before the amendment of the classification and the use of genetic tools to improve the phylogeny. Because the biosynthetic pathway to DGTS is now known, new phylogenetic inference on DGTS biosynthesis enzymes could be achieved (Bolik et al., 2023; Murakami et al., 2018), validating the absence of DGTS synthesis enzyme in the genome of seed plants.

Biosynthesis of DGTS was first discovered in bacteria and is realized in two steps by the two enzymes BtaA and BtaB whereas in *Chlamydomonas* and in fungi it is catalyzed by one bifunctional enzyme Bta1 containing the two domains BtaA and BtaB. BtaA uses diacylglycerol (DAG) and S-adenosylmethionine (SAM) as substrates to produce diacylglycerol-O-homoserine that will then be used by BtaB to produce DGTS by adding successively three methyl groups on the nitrogen from the SAM donor (Riekhof et al., 2005a, 2005b). To investigate the role of betaine lipid, we produced DGTS in plants either by producing transgenic Arabidopsis line expressing Bta1 under the control of a phosphate starvation inducible promoter, pMGD3 (Shimojima et al., 2015), or by overexpressing transiently Bta1 under 35S promoter in *Nicotiana benthamiana*. Our results indicate that DGTS is produced from neosynthesis of lipid and not from lipid remodeling, generating an endoplasmic reticulum (ER) proliferation. DGTS production do not seem to improve growth and to affect lipid remodeling in phosphate starvation.

## Results

### Selection of DGTS synthase gene for DGTS plant production

BTA1 genes from different species were identified by blast using the protein sequence of *Chlamydomonas reinhardtii* BTA1. To identify Bta1 enzymes that are functional in higher plants, five BTA1 sequences from different lineages were selected for functional tests by transient expression in *Nicotiana benthamiana* . Therefore, BTA1 cDNA from *Microchloropsis gaditana* (MgBTA1, eustigmatophyte)*, Phaeodactylum tricornutum (nf-PtBTA1,* diatom)*, Kluveromyces lactis* (KlBTA1, fungi)*, Marchantia polymorpha* (MpBTA1, bryophyte) and *Chlamydomonas reinhardtii* (CrBTA1, green algae) were cloned under the control of 35S promoter and were transiently overexpressed in *N. benthamiana* leaves by agroinfiltration. Eight days after the infiltration, lipids were extracted from the leaves and separated by thin layer chromatography (TLC) (Figure 1). A lipid spot corresponding to DGTS was clearly identified when the enzyme of *M. gaditana, K. lactis, M. polymorpha and C. reinhardtii* were expressed, showing that DGTS could be produced in higher plants by expressing a BTA1 gene. These results also confirmed that the BTA1 gene of *M. polymorpha* and *M. gaditana* encodes functional enzymes. However, we did not detected DGTS production in leaves expressing the BTA1 construct from *P. tricornutum*. By further analyzing the annotated sequence retrieved from the Joint Genome Institute (JGI), we noticed that an intron, not initially annotated, was present in the N-terminus sequenced that was cloned from the genomic DNA of *P. tricornutum* (Figure S1B). We therefore cloned the correct version of Phaeodactylum gene from cDNA and to be sure that the absence of DGTS synthesis was not linked to an ER addressing default, we fused both *P. tricornutum* and *M. gaditana* to the N-terminal ER transmembranes domain of Sec63. We then expressed both constructs in *N. benthamiana* leaves. To better separate DGTS from endogenous lipids, we performed 2D-TLC with lipids extracted from non-transformed plants or plants transformed with the new *P. tricornutum* BTA1 construction (thereafter named PtBTA1) and MgBTA1 as a positive control (Figure S2). This experiment showed that PtBTA1 was indeed functional, but produced DGTS at a lower level in *N. benthamiana* compared to MgBTA1, probably linked to a difference of level of expression of the two proteins (see below). Thus, the presence of the N-terminus intron leads to the production of a non-functional enzyme that we named nf-PtBTA1. For further experiments, MgBTA1, PtBTA1, and nf-PtBTA1 were chosen for a strong, moderate and inefficient production of DGTS in plant, respectively.

**Figure 1.**
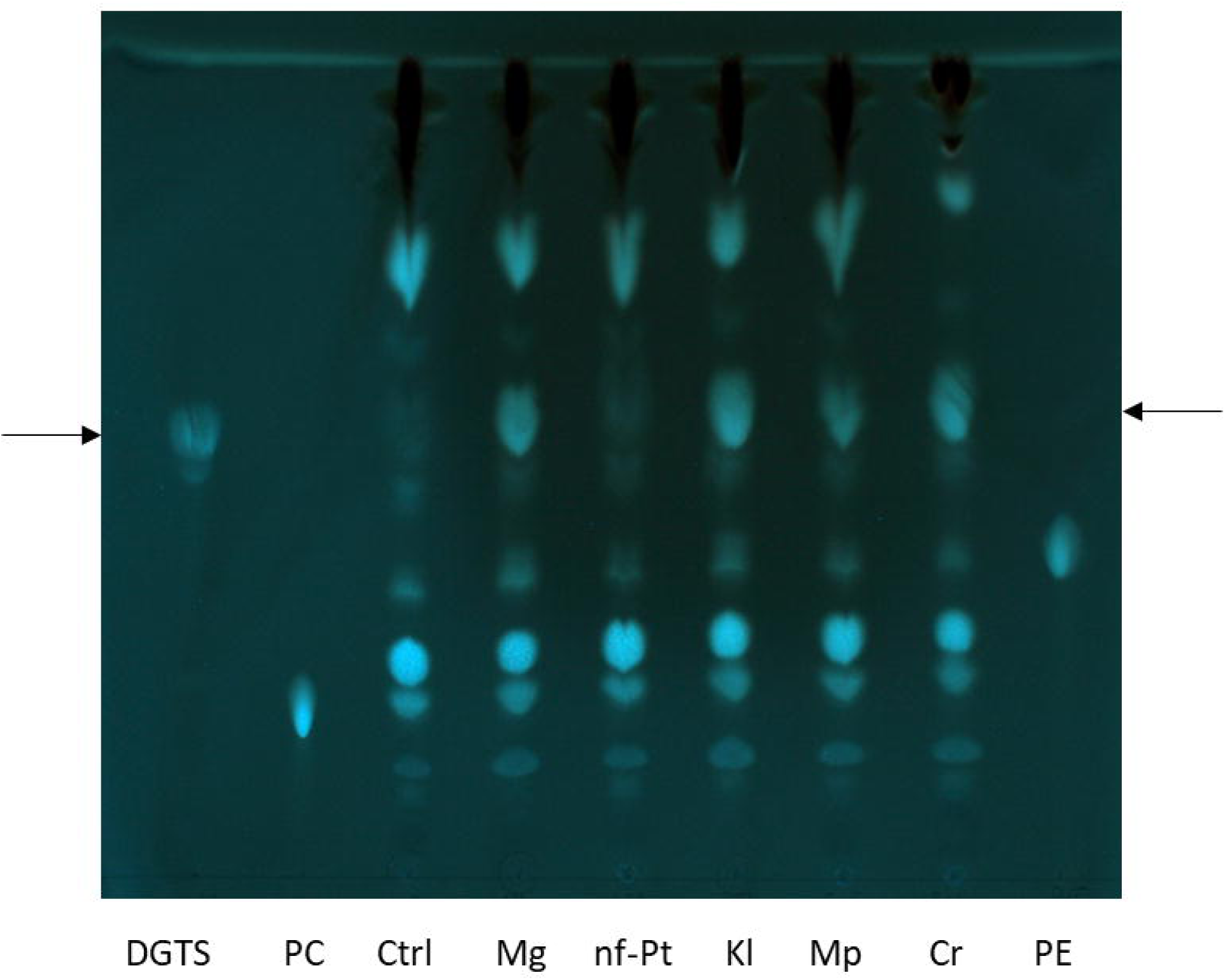
TLC analysis of lipid extracted from. Nicotiana benthamiana **leaves expressing different BTA1 genes.** Lipid control for migration: DGTS, PC and PE. The arrows indicate the DGTS position. Ctrl corresponds to leaf extract without Bta1, all other lines (Mg, nf-Pt, Kl, Mp, Cr) correspond to leaf extract expressing different genes of Bta1. Mg: *Microchloropsis gaditana*; nf-Pt: non functional *Phaeodactylum tricornutum*; Kl: *Kluveromyces lactis*; Mp: *Marchantia polymorpha*; Cr: *Chlamydomonas reinhardtii*.

### Analysis of Arabidopsis seedlings producing DGTS in phosphate starvation

We first tried to transform Arabidopsis with *Mg*BTA1 under the control of the 35S promoter but in our hands, we never managed to obtain transformed seeds. We decided then to express *Mg*BTA1 under a phosphate starvation inducible promoter, pMGD3 (Figure S1C) (Shimojima et al., 2015). We used the construction targeted to the ER by the addition of the transmembrane fragment of Sec63 to ensure that the production of DGTS is localized in the ER. We selected two independent lines that express *Mg*BTA1 during phosphate starvation (Figure S3) and carried on their analysis on seedlings and calli culture.

Col0, pMGD3-MgBTA1-1 and pMGD3-MgBTA1-2 seeds were grown for 21 days *in vitro* on MS media with or without Pi (Figure 2A). As expected, Pi starvation severely slows down plant growth but no major phenotypic difference was detected between the Col0 and lines expressing BTA1 in both conditions. Lipids were extracted from the rosette leaves and analysed by HPLC-MS/MS (Figure 2B and C). In +Pi conditions, lipid content is similar between Col0 and the BTA1 overexpressors (Figure 2B) and no significant difference in lipid composition were seen between Col0 and the mutants (Figure 2C). Furthermore, DGTS was barely detected in the mutants confirming that genes under the control of pMGD3 are not expressed in +Pi condition. In –Pi conditions, the plant lipid content is slightly decreased compared to +Pi condition, but no significant difference between Col0 and the mutants is observed (Figure 2B). DGTS synthesis is induced in the mutants reaching up to 4 % of total glycerolipids but the rest of the glycerolipidome is not significantly affected. Phospholipids are degraded and DGDG production is significantly increased by Pi starvation as previously observed (Figure 2C). DGTS production does not seem to affect glycerolipid remodeling, probably because the induction of DGTS synthesis is not very strong.

**Figure 2.**
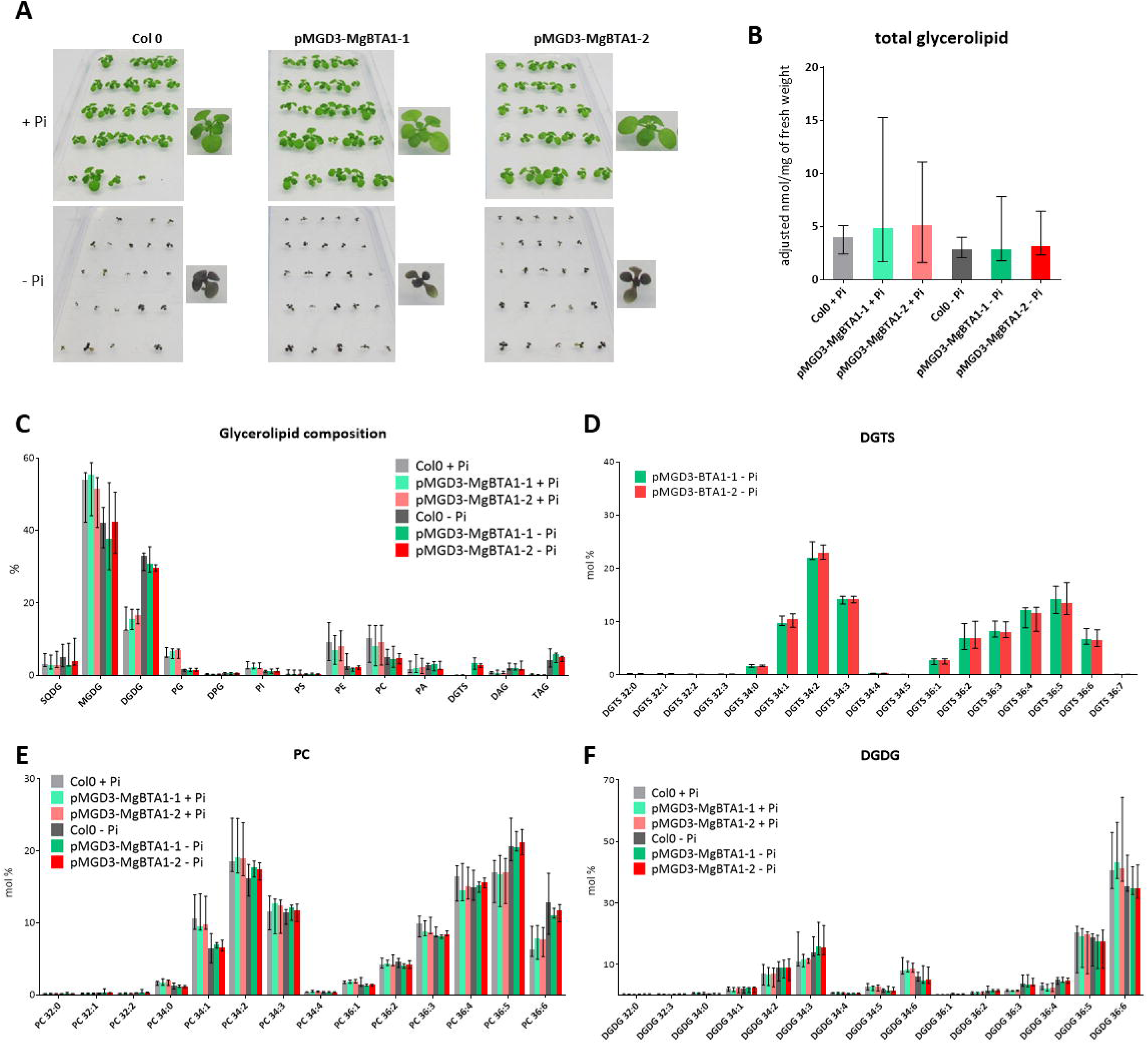
pMGD3-BTA1Mg-HA **seedlings analysis. (A)** Pictures of *pMGD3-BTA1Mg-HA* (203-2-5 n°1 and 203-4-1 n°3 lines) and Col0 seedlings grown during 21 days on MS medium supplemented with 1 mM Pi (+Pi) or 0,005 mM Pi (-Pi). **(B)** to **(F)** Lipid analysis of *pMGD3-BTA1Mg-HA* (203-2-5 n°1 and 203-4-1 n°3 lines) and Col0 seedlings leaves grown during 21 days on MS medium supplemented with 1 mM Pi (+Pi) or 0,005 mM Pi (-Pi). Represented values correspond to the median of 5 biological replicates and the error bars represent the range between minimum and maximum values. **(B)** Quantity of glycerolipids in nmol per mg of rosette fresh weight. **(C) (D)**, **(E)**, **(F)** Glycerolipid distribution in mol % **(C)** and molecular distribution within DGTS (D), PC (E) and DGDG (F) classes determined by HPLC-MS/MS. The first number corresponds to the sum of the carbon numbers of the two fatty acids, and the second number to the number of unsaturations. SQDG, sulfoquinovosyldiacylglycerol; MGDG, monogalactosyldiacylglycerol; DGDG, digalactosyldiacylglycerol; PG, phosphatidylglycerol; DPG, diphosphoglycerate; PI, phosphatidylinositol; PS, phosphatidylserine; PE, phosphatidylethanolamine; PC, phosphatidylcholine; PA, phosphatidic acid; DGTS, ; DAG, diacylglycerol; TAG, triacylglycerol.

We also analysed the distribution of DGTS molecules in –Pi, as DGTS was barely detectable in +Pi condition (Figure 2D). Each molecule is described first by the number of carbon in the acyl groups and secondly by the number of unsaturations. For example, DGTS 36:2 is a 1,2-diacyl-*sn*-glycero-3-trimethylhomoserine in which the acyl groups at *sn*-1 and *sn*-2 positions contain 36 carbons in total and 2 double bonds. DGTS molecule distribution and desaturation pattern are similar to those of PC molecules (Figure 2E) but with a slighter enrichment of 34 C diacylglycerol backbone, corresponding to a C16 and a C18 fatty acid, compared to PC enriched in 36 C diacylglycerol backbone, corresponding to two C18 fatty acids. Because *Microchloropsis* is an algae containing very few fatty acids with C18 (Billey et al., 2021), this difference could be due to the enzyme specificity. Because chloroplast lipids (MGDG, DGDG, SQDG and PG) have peculiar lipid compositions (Figure 2F and Figure S4A-C) distinct from DGTS and because DGTS composition is close to PC composition, DGTS is probably produced in the ER by the Kennedy pathway and thus uses the same DAG pool as PC (Karki et al., 2019).

### Analysis of Arabidopsis calli producing DGTS in phosphate starvation

To confirm this hypothesis and to see if DGTS production has an impact on the kinetic of lipid remodeling in phosphate starvation, calli were produced from the same Col0, pMGD3-MgBTA1-1 and pMGD3-MgBTA1-2 and lines and grown in replete condition (+Pi) and in starved condition (-Pi) for 4 and 8 days (Figure 3). Lipids were extracted and analysed by HPLC-MS/MS. Calli are non-photosynthetic cells and therefore the ratio of phospholipids over galactolipids is much higher than in leaves and lipid remodeling defects can be more visible in calli. Phosphate starvation triggers in 8 days a small, but non-significant, diminution of the total glycerolipid content (Figure 3A), phospholipids are degraded and DGDG and SQDG increased (Figure 3B). In the transformant lines, DGTS increased more than 100 times (Figure 3C) but represents less than 5 % of the total glycerolipid content, even after 8 days of starvation. Similar to what we observed in leaves, the synthesis of DGTS had no major influence on the kinetic of lipid remodeling, except maybe a small non-significant delay in the decrease of the glycerolipid content that could be explained by the neosynthesis of DGTS (Figure 3C).

**Figure 3.**
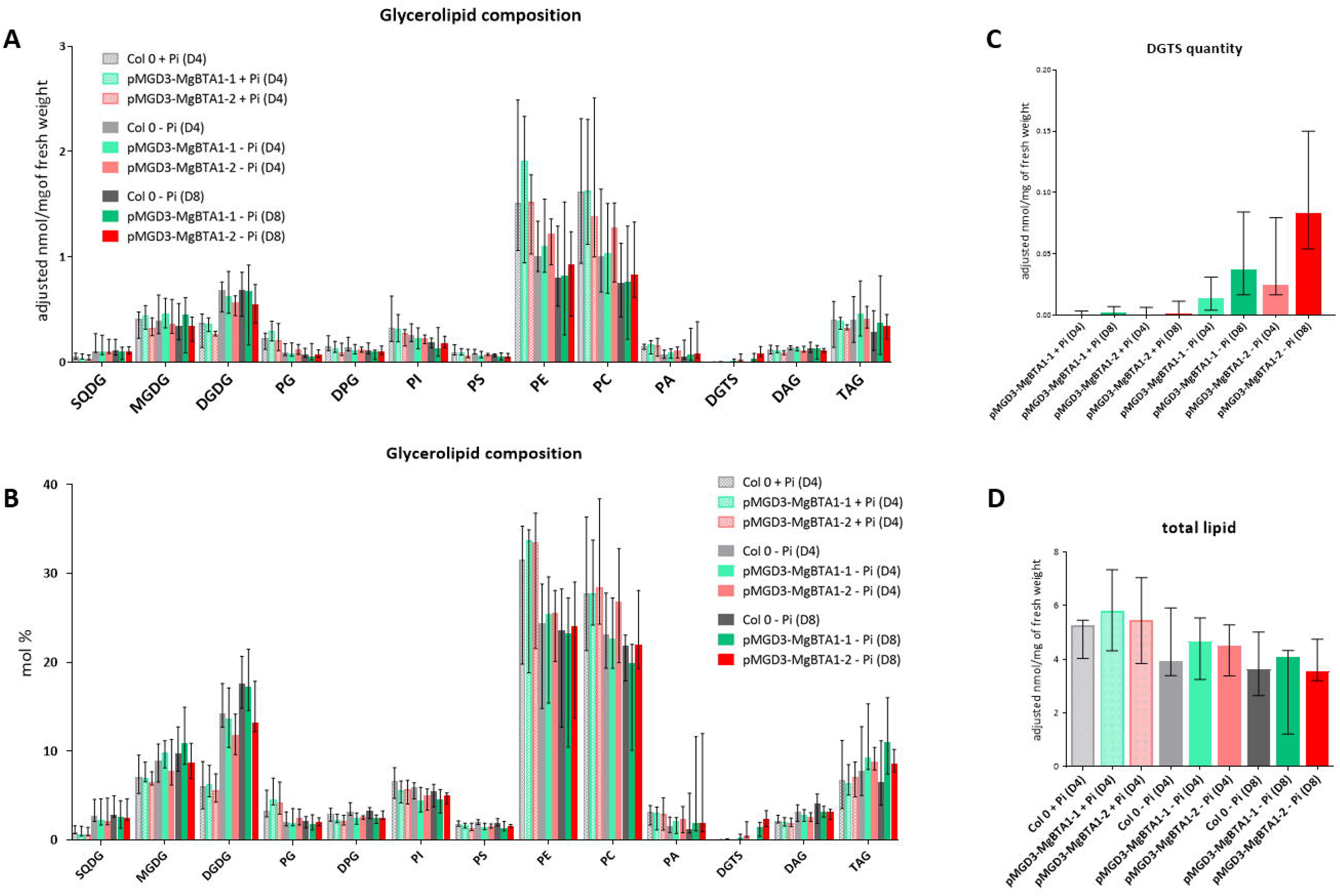
Lipid analysis of. pMGD3-BTA1Mg-HA **calli. (A)** to **(D)** Lipid analysis of *pMGD3-BTA1Mg-HA* and Col0 whole calli grown in phosphate replete (+Pi) and deplete (-Pi) conditions and harvested at day 4 (D4) and day 8 (D8). Represented values correspond to the median of 6 biological replicates and the error bars represent the range between minimum and maximum values. **(A)** Adjusted glycerolipid content per mg of fresh weight, **(B)** glycerolipid composition in mol %, **(C)** zoom on DGTS content and **(D)** total lipid content, all determined by HPLC-MS/MS. SQDG, sulfoquinovosyldiacylglycerol; MGDG, monogalactosyldiacylglycerol; DGDG, digalactosyldiacylglycerol; PG, phosphatidylglycerol; DPG, diphosphoglycerate; PI, phosphatidylinositol; PS, phosphatidylserine; PE, phosphatidylethanolamine; PC, phosphatidylcholine; PA, phosphatidic acid; DGTS, ; DAG, diacylglycerol; TAG, triacylglycerol.

When analyzing fatty acid composition and glycerolipid molecules distribution (Figure S5A), mutant cells are enriched in 18:1 fatty acid. This phenomenon is visible in +Pi condition and is therefore independent of DGTS production. It is more probably due to a difference of the kinetic of cell division as previously observed and not to the mutation (Meï et al., 2015). This difference is reflected in every glycerolipid composition (Figure S5C and D). Looking at the DGTS composition (Figure S5B), as in leaves, DGTS molecule distribution is similar to PC with a slight enrichment in species with 34C and very different than the composition of DGDG, reflecting DGTS is synthesized from a distinct pool of DAG dedicated to extra-plastidial lipid synthesis.

### Analysis of Nicotiana benthamiana leaves overproducing DGTS

Because DGTS production in our stable *A. thaliana* lines was very low, we used the transient expression system in *N. benthamiana* described above to obtain a significantly higher level of DGTS synthesis and investigate the impact of this lipid on higher plant lipidome and membranes structure. Overexpression of protein by agroinfiltration has an impact on leaf metabolism including shutdown of chloroplast gene expression and activation of oxidative stress responses (Hamel et al., 2024). To be sure that the phenotypes observed on leaf are due to DGTS production and not to protein overexpression and agroinfiltration stress, three DGTS synthases targeted to the ER with different amount of DGTS production were chosen to be overexpressed: BTA1 from MgBTA1, for a strong DGTS production, PtBTA1 for a medium production, and nf-PtBTA1 for ineffective production (Figure S1B).

The three enzymes were fused to GFP in N-terminus to validate their localizations and the GFP fluorescence was observed in confocal microscopy (Figure 4). For all three enzymes, the GFP signal resembled ER membrane pattern in epidermis cell with the standard reticulated network pattern and surrounding of the nucleus (Figure 4A). As variations in the GFP signal intensities were reproducibly observed between the three constructions, we quantified the level of GFP fluorescence within cells from independent experiments. The fluorescence intensity level of MgBTA1-GFP and nf-PtBTA1-GFP was consistently higher than the one measured for PtBTA1 (at least twice higher), indicating a lower expression of the enzyme (Figure 4B). These results can explain the lower DGTS production observed by TLC (Figure S2). To confirm this results, we tried to quantify the BTA1-GFP signal by Western blot however, whatever the protein extraction method and the protease inhibitors used, no signal corresponding to the intact protein was detected but a signal corresponding to the GFP alone was measured (Figure S6). Because the fluorescence signal is not cytosolic as expected for GFP alone, *in planta*, the BTA1-GFP fusion should be preserved. The intensity of the GFP band is related to the level of fluorescence observed, very high for MgBTA1-GFP and nf-PtBTA1-GFP and barely detectable for PtBTA1-GFP. To validate the ER localization, MgBTA1-GFP and nf-PtBTA1-GFP were co-expressed with an ER marker fused to RFP (Figure 4C). GFP and RFP are co-localized, indicating that our construct localization is well restricted to the ER and that probably no degradation of the fusion releasing GFP alone occurs *in planta*.

**Figure 4.**
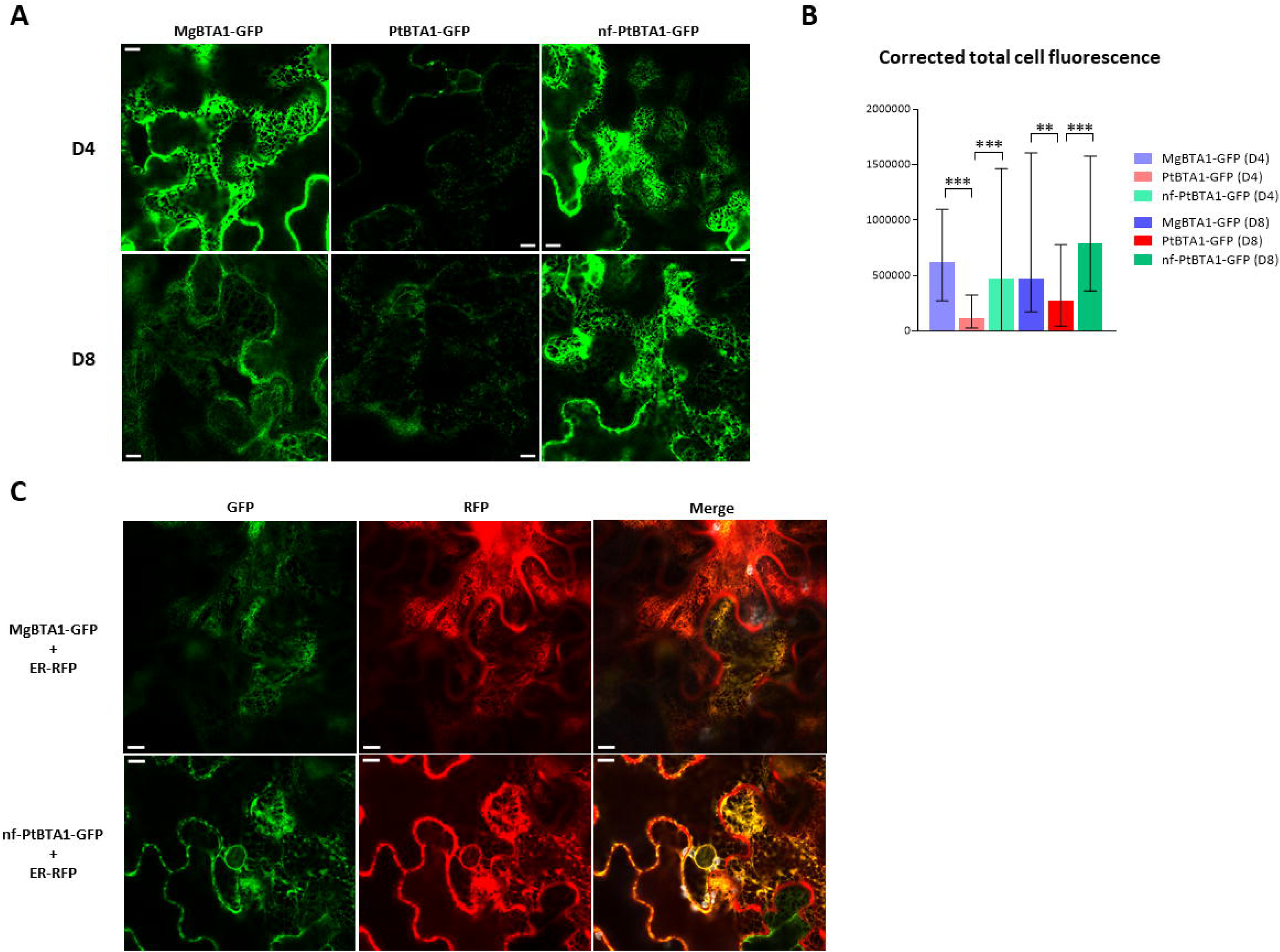
**Subcellular localizations of BTA1 proteins onverexpressed in**Nicotiana benthamiana**. (A)** Subcellular localization of MgBTA1-GFP, PtBTA1-GFP and nf-PtBTA1-GFP proteins transiently expressed in *N. benthamiana* leaves 8 days after infiltration. (Scale bar, 10 μm). **(B)** Fluorescence quantification from confocal images of BTA1-GFP, PtBTA1-GFP and nf-PtBTA1-GFP fused proteins transiently expressed in *N. benthamiana* leaves 8 days after infiltration. Determination of Corrected total cell fluorescence (CTCF) was done with ImageJ software with a total of 30 images per construction, taken from 3 independent plants. CTCF = Integrated Density – (Area of selected cell X Mean fluorescence of background readings). Asterisks indicate significant differences (\*\**P*<0,01*;***P*<0,001) in pairwise comparisons by a Kruskal-Wallis test. **(C)** Subcellular localization of MgBTA1- GFP and nf-PtBTA1-GFP proteins transiently co-expressed with the ER marker ER-RFP in *N. benthamiana* leaves. ER-RFP : RFP protein fused to a HDEL retention signal in C-ter. (Scale bar, 10 μm).

Lipids from leaves transfected with P19 only (infiltration control), nf-PtBTA1-GFP, PtBTA1-GFP and MgBTA1- GFP fused proteins, 4 and 8 days after agroinfiltration, were analyzed by HPLC-MS/MS (Figure 5 and S7). Surprisingly the construct nf-PtBTA1-GFP was able to produce a very small amount of DGTS, not detectable on the TLC (Figure 1), indicating that the function of the enzyme with an extended N-terminus is not completely impaired (Figure 5A). However, the synthesis of DGTS is negligible in comparison to the leaves expressing PtBTA1-GFP and MgBTA1-GFP, despite a protein expression similar to MgBTA1 (Figure 4B). In PtBTA1 and MgBTA1 overexpressors, DGTS accumulates at a very high level, even equivalent to MGDG content at D8 in the MgBTA1 overexpressing leaves (Figure 5A and B), reflecting probably the difference of expression of the two proteins. Indeed, DGTS represented up to 30 mol% of glycerolipid content in leaves overexpressing MgBTA1- GFP (Figure 5B). When looking at the DGTS composition, as in Arabidopsis, DGTS molecule distribution is comparable to PC distribution with a preference of DAG backbone of 34 carbons and is very distinct from DGDG distribution (Figure S7A-C). This enrichment of 34 carbons in DGTS can be seen at the level of the fatty acid composition in MgBTA1 agroinfiltrated leaves with a global enrichment of 16:0 (Figure S7D).

**Figure 5.**
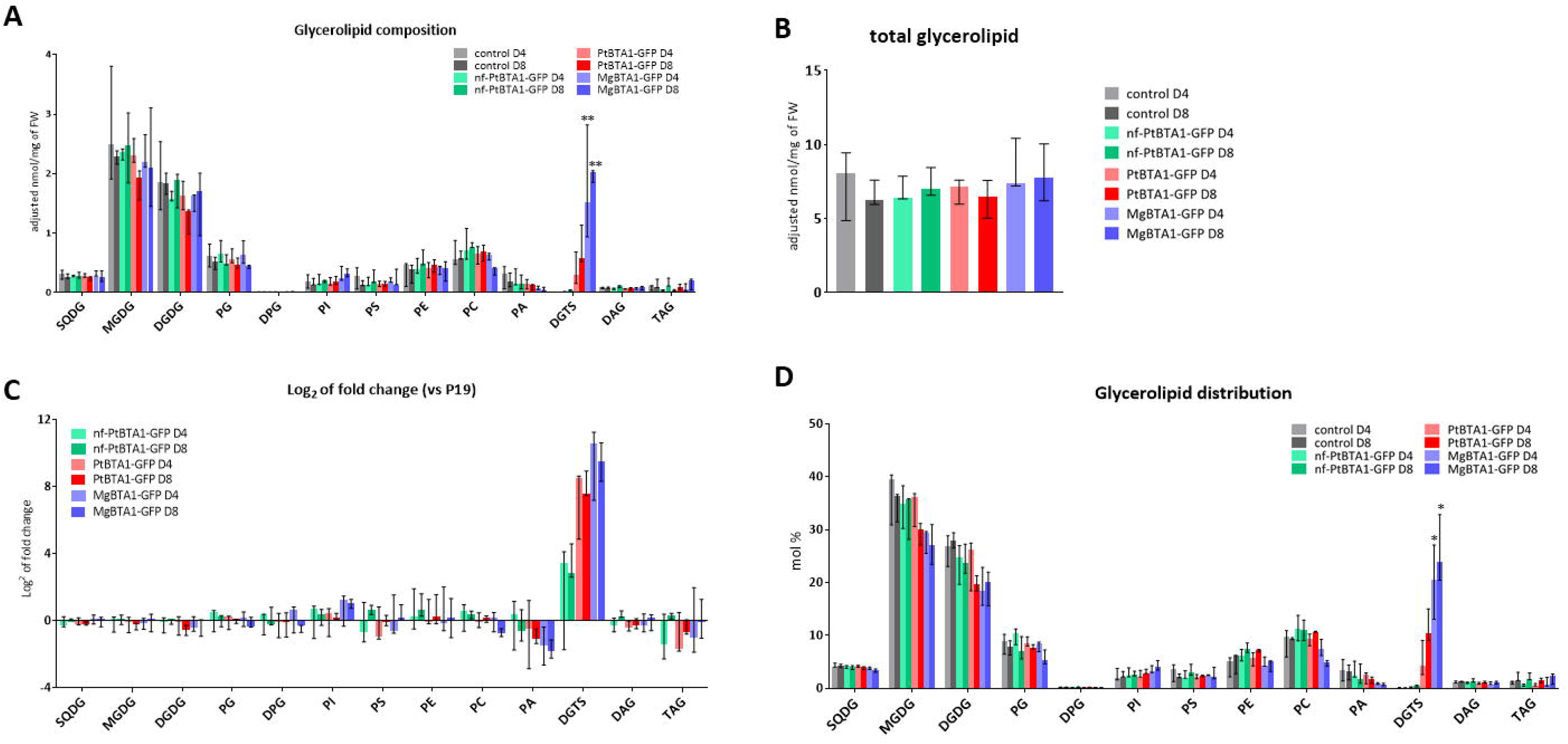
Lipid analysis of. Nicotiana benthamiana **leaves agroinfiltrated with BTA1 constructions. (A)** to **(D)** Lipid analysis of *N. benthamiana* leaves agroinfiltrated with P19 only, nf-PtBTA1-GFP, PtBTA1-GFP and MgBTA1-GFP fused proteins. Leaves were harvested 4 (D4) and 8 (D8) days after agroinfiltration. Control: *N. benthamiana* agroinfiltrated with P19 protein (negative control). Represented values correspond to the median of 3 biological replicates and the error bars represent the range between minimum and maximum values. Asterisks indicate significant differences (\*\**P*<0,01 ; \**P*<0,05) in comparison to control D4 and D8 by a Kruskal-Wallis test. **(A)** Quantity of each class of glycerolipids determined by HPLC-MS/MS. **(B)** Total glycerolipid quantity determined by HPLC-MS/MS. **(C)** Log_2_ of fold change in lipid quantities between BTA1 constructions (nf-PtBTA1-GFP, PtBTA1-GFP and MgBTA1-GFP) and P19.**(D)** Glycerolipid distribution. SQDG, sulfoquinovosyldiacylglycerol; MGDG, monogalactosyldiacylglycerol; DGDG, digalactosyldiacylglycerol; PG, phosphatidylglycerol; DPG, diphosphoglycerate; PI, phosphatidylinositol; PS, phosphatidylserine; PE, phosphatidylethanolamine; PC, phosphatidylcholine; PA, phosphatidic acid; DGTS, ; DAG, diacylglycerol; TAG, triacylglycerol.

However, this high amount of DGTS does not have a huge impact on global glycerolipid content, except maybe a slight non-significant increase of lipid content in MgBTA1 overexpressor (Figure 5C). Looking at the glycerolipid proportion within the leaves (Figure 5B) although the difference are non-significant, galactolipids seem to be affected when DGTS is produced above 10 % of the glycerolipid content whereas SQDG is not. In MgBTA1 overexpressor, PA and PC content seems also to decrease as well as PG at D8 to compensate DGTS increase.

Due to the huge amount of DGTS production in the MgBTA1 overexpressor and the absence of major impact on other glycerolipids content, we were wondering where the produced DGTS molecules were localized in cells, particularly regarding chloroplast membranes, representing more than 70 % of the cell membranes. Chloroplast were purified from D4 agroinfiltrated leaves with nf-PtBTA1-GFP, as a protein overexpressor and poor producer of DGTS control, or with MgBTA1-GFP leaves, that contain more than 20 % of DGTS (Figure 6A and B). Chloroplast fractions are characterized by a high proportion of glycolipid and a very low amount of PE and PC in comparison to the total fraction whereas the opposite is true for endomembrane fraction. The chloroplast fractions for both overexpressors are pure, with an enrichment of LHCP (thylakoid marker) and no labelling of BIP2 and GFP (Figure 6C). In these fractions, DGTS was detected at the same level in nf-PtBTA1 and MgBTA1 agroinfiltrated leaves, less than 1.5 % (Figure 6B). The extraplastidial fractions are contaminated by chloroplast with a high amount of galactolipid and the presence of LHCP, however they are enriched in endomembranes with strong BIP2 signal and an enrichment in PE absent from chloroplast membrane. This fraction contains roughly the same amount of DGTS than the total extract. Overall, this experiment indicate that DGTS is absent from chloroplast and restricted to extraplastidial membranes, highlighting the specificity of lipid transport pathway to chloroplasts.

**Figure 6.**
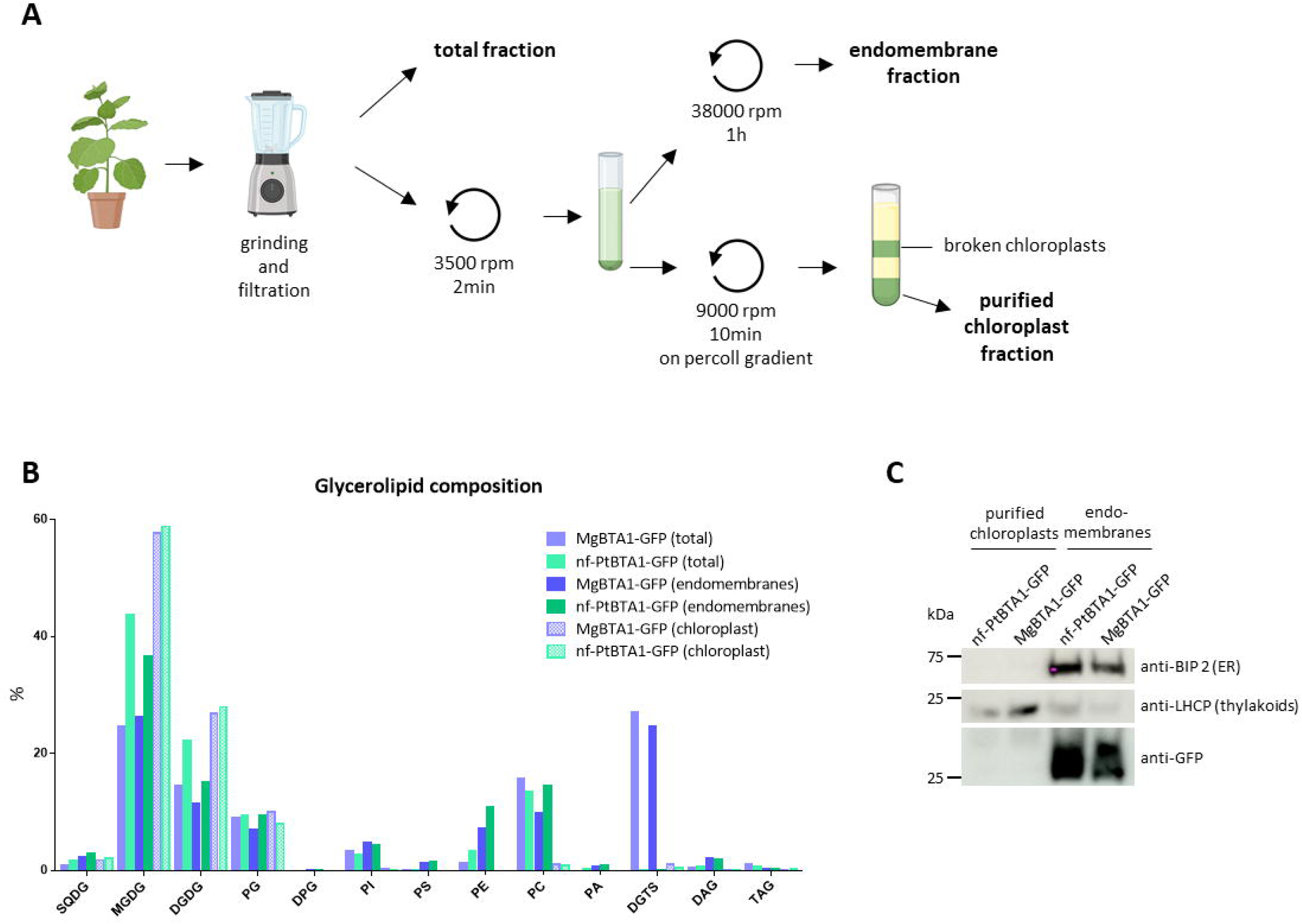
Localisation of DGTS in. Nicotiana benthamiana **leaves overexpressing either MgBTA1-GFP or nf- PtBTA1-GFP. (A)** Strategy used to isolate chloroplast and extraplastidial fractions from *N. benthamiana* leaves 4 days after agroinfiltration with BTA1Mg-GFP and nf-BTA1Pt-GFP fused proteins. Curved arrows represent the centrifugation steps. **(B)** Proportion of glycerolipids determined by HPLC-MS/MS in each fraction. Total, total fraction; extraplast mb,extraplastidial membrane fraction; chloroplast, purified chloroplasts fraction. **(C)** Western blot analysis of purified chloroplasts and extraplastidial membranes fractions using BIP 2 antibody for ER, LHCP for thylakoids and GFP for BTA1 fused proteins.

To measure the impact of DGTS accumulation on extraplastidial membranes, leaves agroinfiltrated at D4 with P19, nf-PtBTA1-GFP, PtBTA1-GFP and MgBTA1-GFP were observed by electronic microscopy (Figure 7A and S8). Chloroplast and thylakoid ultrastructure looks unaffected, except maybe an increase in starch content in MgBTA1 overexpressor. To validate if it was not an artefact of observation, starch content was measured in different leaf extract from different independent plants and leaves. The starch content was highly variable depending on the experiment and seems to be more related to the level of protein overexpression than DGTS production, nf-PtBTA1 and MgBTA1 having the highest level of protein expression and starch content with very different DGTS amount (Figure S9A). Photosynthesis efficiency was also measured (Figure S9B and C). As expected, agroinfiltration has a negative impact on the electron transfer rate (ETR) and maybe a slight delay on the relaxation time of the non photochemical quenching (NPQ) (Hamel et al., 2024). No difference was observed between P19 and the BTA1 overexpressor in terms of NPQ. However, the ETR seems to be slightly related to the DGTS amount with an ETR value following the DGTS proportion: ETR_MgBTA1_ ≤ ETR_PtBTA1_ ≤ ETR_nf-PtBTA1_ and ETR_P19_. Further investigations would be needed to confirm if this finding is related to the observed galactolipid decrease.

**Figure 7.**
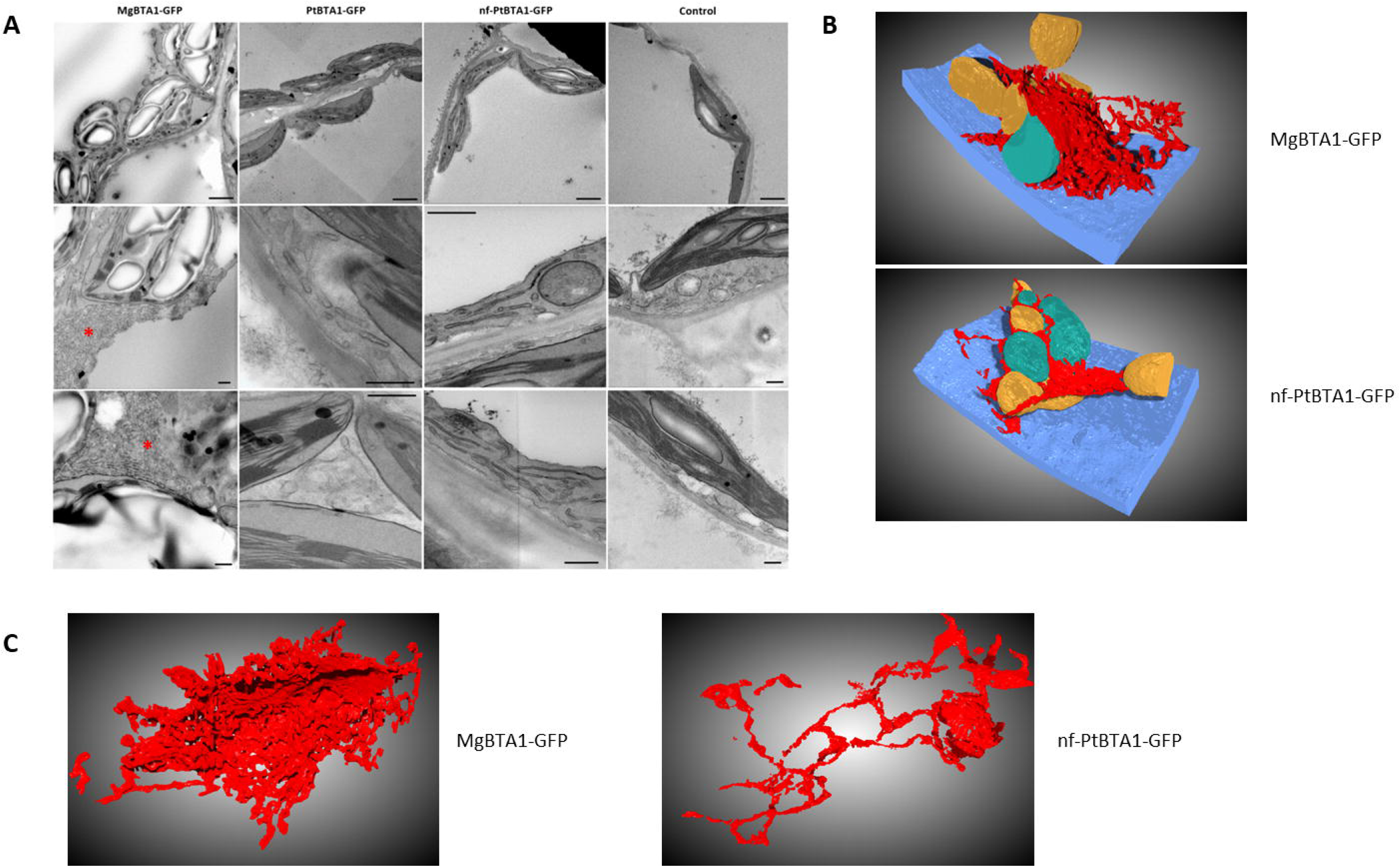
Impact of BTA1 proteins transient expression on. Nicotiana benthamiana **leaves cell architecture (A)** Electronic microscopy observation of the architecture of cellular membranes in *N. benthamiana* leaves agroinfiltrated with BTA1 fused proteins (nf-PtBTA1-GFP, PtBTA1-GFP and MgBTA1-GFP) or P19 as control at day 4. The scale bar is 2 µm for the top four images and 500 nm for the rest of the images. Red asterisks indicate membrane ER proliferation. **(B)** and **(C)** Representative 3D architecture of the cell (B) and the ER alone from FIB-SEM imaging from *N. benthamiana* leaves agroinfiltrated with MgBTA1-GFP or nf-PtBTA1-GFP. In blue, the cell wall; in red, the ER; in yellow, the mitochondria and in green the peroxisomes. These pictures are extracted respectively from the supplementary movies 1 for MgBTA1-GFP and 2 for nf-PtBTA1-GFP.

In MgBTA1 overexpressor, membrane proliferation was observed in the cytosol that could be related to the extraplastidial localization of DGTS (Figure 7A and S8). The architecture of these membrane structures was looked in 3D by Focused Ion Beam Scanning Electron Microscopy (FIB-SEM) in comparison with what could be observed in nf-PtBTA1 overexpressor (Supplemental movie 1 and 2). A snapshot of the reconstituted cell architecture showing the cell wall, the mitochondria, the peroxisome and the ER without the plastid is presented Figure 7B. In both case, ER is surrounding mitochondria and peroxisomes that are embedded within ER tubules. Furthermore, reconstructed ER from MgBTA1-GFP overexpressor is constituted of numerous tubule layers but with a membrane organization very similar to the standard ER architecture observed in nf-PtBTA1 overexpressor (Figure 7C). In reference to the cell wall volume, the volume of the ER in MgBTA1-GFP overexpressor is 8 times bigger than the volume of the ER in nf-PtBTA1-GFP. Overall, these results indicate that DGTS is probably synthesized in the ER and its accumulation stimulates ER proliferation at the expense of the other membranes.

## Discussion

Betaine lipids are absent in seed plants but could be found in chlorophytes up to ferns (Rozentsvet, 2004; Vogel and Eichenberger, 1992). Since the identification of the genes responsible for DGTS synthesis (Riekhof et al., 2005a, 2005b), genome search could be achieved as well as phylogenetic reconstruction that confirm the absence of BTA1 gene in seed plants, gymnosperm and angiosperm, and the evolution of the gene link to the habitat, fresh water *versus* marine environment (Bolik et al., 2023). By expressing BTA1 genes derived from marine or fresh water organisms in *Nicotiana benthamiana* (Figure 1), we demonstrated that plants are still able to produce DGTS and that only the absence of the BTA1 gene is responsible for the loss of their capacity to synthetize DGTS. However, in our hands, we never managed to obtain stable lines of *Arabidopsis thaliana* that were overexpressing DGTS under the control of the constitutive promoter 35S. Previous work indicates that DGTS bilayer are thicker than PC bilayer and that the thickness could be sensitive to hydration (Bolik et al., 2023). Seed plants lost their desiccation tolerance in vegetative stage compared to primitive plant and produced seeds representing a very dry stage (less than 20 % of protoplasmic water) (Oliver et al., 2000). The disappearance of BTA1 gene is concomitant to these two parameters. Further experiments would be needed to see if it is just a correlation or a causal link.

Betaine lipids are known to substitute to extraplastidial phospholipids in different organisms, bacteria, fungi and algae in phosphate starvation (Abida et al., 2015; Mühlroth et al., 2017; Murakami et al., 2018; Oishi et al., 2022; Riekhof et al., 2014; Van Mooy et al., 2009) whereas in seed plants, this function is assured by DGDG that will be synthesized in plastids and exported outside of the chloroplast (Andersson et al., 2005; Jouhet et al., 2004). We decided to stably express in *Arabidopsis thaliana* MgBTA1 under the control of phosphate starvation inducible promoter, pMGD3 (Shimojima et al., 2015). In phosphate starvation condition, DGTS was able to be synthesized in vegetative tissues at a very low level without having any impact on plant growth or lipid remodeling. DGDG content increased and phospholipids degradation was similar to the wild type plants. The level of DGTS production was quite low, less than 5 % of total glycerolipids. pMGD3 was shown to be induced at least 10 times in phosphate starved condition compared to control condition (Shimojima et al., 2015) whereas 35S promoter is supposed to induce the expression more than 100 times for the same genes (Demski et al., 2020). The difference of promoter strength is likely the explanation for the low DGTS production. Indeed DGTS synthesis occurs in two steps: the first one uses DAG and SAM as substrates to produce diacylglycerol-O-homoserine that will then be used to produce DGTS by adding successively three methyl groups on the nitrogen from the SAM donor (Riekhof et al., 2005a, 2005b). During phosphate starvation, DAG is supposed to be released from PC to be recycled into DGDG (Jouhet et al., 2003; Nakamura et al., 2009), however the localization of DAG release is still unknown. Because DGTS synthesis in phosphate starvation does not impact DGDG synthesis neither affect its molecular composition (Figure 2, 3 and S5), DAG produced from PC degradation is not accessible to BTA1 enzyme, either because this process does not occurred in the ER or because DAG is never freely available and directly channels to chloroplast outer membrane for DGDG synthesis. Nevertheless, the low level of DGTS obtained even after 8 days of starvation could explain the absence of phenotypes in our stable lines and the use of a promotor with a higher expression level is required to clearly conclude about the role of DGTS in lipid remodeling and plant acclimation to phosphate starvation.

In *N. benthamiana* leaves, when using 35S promoter, DGTS was able to accumulate up to 20% of total glycerolipid at a slight expense of galactolipid, PC and PA production (Figure 5). However, the molecular distribution of all these lipid classes did not change (Figure S7), showing that even a high level of DGTS does not affect the eukaryotic pathway in galactolipid synthesis (Petroutsos et al., 2014) and therefore is not interfering with the recycling of DAG backbone coming from PC into galactolipids. DGTS synthesis seems therefore to occur more from DAG *de novo* synthesis, thus from the Kennedy pathway, than to derive from PC backbone (Zhou et al., 2020). If this hypothesis is true, it means fatty acid desaturases are able to work on DGTS as well as on PC. In *C. reinhardtii*, that does not contain PC, fatty acid desaturases are able to work on DGTS (Li-Beisson et al., 2015) and CrFAD2 sequence is homolog of AtFAD2 (Chi et al., 2008), making this hypothesis plausible and suggesting that the ability to desaturate DGTS was not lost through the evolution. Furthermore, a very high amount of DGTS stimulates slightly fatty acid production with an enrichment in 16:0 fatty acid (Figure S7D). This is consistent with the composition of DGTS enriched in DAG backbone with 34 carbon - i.e. 16:0/18:X - in comparison with PC composition and could be related to a “pull” effect (Vanhercke et al., 2019) generated by the consumption of *de novo* DAG by BTA1 that will drag indirectly fatty acid synthesis. Another possibility for the accumulation of DGTS would be a “protect” effect (Vanhercke et al., 2019) with the loss of lipases able to degrade betaine lipid during the evolution. Lipases acting on betaine lipid to produce DAG have not been identified yet (Hoffmann and Shachar-Hill, 2023) and phospholipase D and C are acting around the phosphate moiety that is not conserved in betaine lipids and therefore are not supposed to be active on this class of lipids. BTA1 gene was targeted to the ER with the Sec63 transmembrane fragment to mimic the localization observed in algae and fungi (Künzler et al., 1997; Zhang et al., 2016). In our experiment, BTA1 was localized in the ER and DGTS accumulation drives an ER proliferation without altering its structure. This feature suggests that DGTS is indeed able to fulfill phospholipid structural role, at least in well-watered leaves. Production of inducible line with hormone treatment such as dexamethasone would be a useful tool to investigate the sensitivity to drought of DGTS overexpressing plant and the *in vivo* dehydration effect on ER membrane enriched in DGTS. The ER proliferation and the absence of DGTS in chloroplast purified fraction indicates that DGTS is produced in the ER and not transported into the chloroplasts. It confirms that lipid trafficking from the ER toward chloroplast membrane is selective as it was already shown that PC was able to be transferred from liposomes to chloroplast but not PE (Yin et al., 2015). The transfer of DGTS to the rest of the endomembrane system could be achieved by vesicular trafficking but this need to be investigated as well as a potential mitochondria transfer. In *C. reinhardtii* and *Saccharomyces cerevisiae* (Li-Beisson et al., 2015; Riekhof et al., 2014), DGTS is replacing PC in every membrane suggesting that DGTS could be present in every extraplastidial membranes.

To resume, DGTS can be synthesized in seed plant leaves and calli by overexpressing BTA1 gene. DGTS synthesis pathway seems to compete with PC synthesis via the Kennedy pathway but does not seem to be derived from PC diacylglycerol backbone and therefore does not interfere with the eukaryotic pathway involved in galactolipid synthesis. DGTS accumulation drives ER membrane proliferation confirming a restricted localization of DGTS in endomembrane and maybe mitochondria and its absence in chloroplast membranes. The use of ER produced DAG as substrate by BTA1 that does not affect galactolipid composition emphasizes our lack of knowledge on the mechanisms and on the nature of the lipid molecules transported to the chloroplast for galactolipid synthesis (Kalisch et al., 2016; Karki et al., 2019; Zhou et al., 2020). Finally, effect of DGTS production needs now to be investigated in drought condition or in seed tissue to see if hydration is indeed a key factor explaining DGTS loss in seed plants.

## Material and Methods

### Plant Materials and Growth Conditions

*Arabidopsis thaliana* ecotype Col0 plants were used as wild-type. Two T-DNA insertion lines in Col0 background were obtained with pMGD3-Sec63TM-BTA1Mg-HAx3 construction (Figure S1) using *BTA1* gene from *Microchloropsis gaditana* (Naga_100016g36). The presence of the T-DNA was verified by PCR using a forward primer (ctggatgggggcccagaagg) located on *BTA1Mg* gene and a reverse primer (agaagcgtagtcttgaacgtcgtatgggtactgggcttttgatttgccgg) located on the junction between *BTA1Mg* gene and HAx3 tag.

Calli were obtained from mesophyll tissue of two week old Arabidopsis leaves, and grown on agar plates containing Murashige and Skoog (MS) (MSP09, Caisson Laboratories, Inc, USA) supplemented with 3 % (w/v) sucrose, 1.2 mg.L^-1^ 2,4-dichlorophenoxyacetic acid and 0.8 % (w/v) agar. After formation, calli were maintained in 200 mL of liquid MS (MS without KH_2_PO_4_, Duchefa Biochemie) supplemented with 0.5 mM of phosphate, 1.5 % (w/v) sucrose and 1.2 mg.L^-1^ 2,4-dichlorophenoxyacetic acid. Calli were kept under continuous light (100 μE.m^-2^.s^-1^) at 22°C, agitated with rotary shaking at 125 rpm and subcultured every 7 days. For phosphate starvation experiments, calli were washed three times with 50 mL of MS medium -Pi (MS without KH_2_PO_4_, Duchefa Biochemie) with 0.5 mM Pi or without Pi and grown for 4 or 8 days in MS with 0.5 mM or without Pi. Arabidopsis mutant and Col0 seedlings lines were cultivated *in vitro* on MS medium (MS without KH_2_PO_4_, Duchefa Biochemie) supplemented with 1 % (w/v) sucrose, 12% (w/v) agar (A7921, Sigma-Aldrich) and 1 mM Pi for Pi replete condition and 0.005 mM Pi for Pi deplete condition. Seedlings were grown 21 days in long day (16h-8h light-dark photoperiod) conditions at 20-22°C with a photon flux density of 30-60 µmol/m^2^/s. On 21- day-old Arabidopsis seedlings, the foliar portion above the hypocotyl was harvested, frozen in liquid nitrogen and stored at -80°C.

*Nicotiana benthamiana* wild type plants were grown in long day conditions (16h-8h light-dark photoperiod) at 18-20°C with 60-70% humidity and a photon flux density of 100-120 µmol/m^2^/s. Plants were cultivated during 4 to 5 weeks on soil (35 L) supplemented with vermiculite (7 L) and biological fungicide (Prestop, Lallemand, 10 g) and watered 3 times a week.

### Arabidopsis transgenic line construction

Genes were cloned in the pFP108 plant expression vector (To et al., 2006) in frame with 3xHA tags added in C-terminal (pFP108-3xHA) and the pMGD3 promoter. Expression of genes from this vector is under the control of a NOS terminator (Figure S1). Plasmids were transferred in *Agrobacterium tumefaciens* C58C1 strain and used from stably transformation of Col0 plants by floral dip. Transformed plants were selected using Basta® and leaves from F2 plants expressing the constructions were used for calli formation.

### Cloning of BTA1 fused proteins used for transient expression in Nicotiana benthamiana

BTA1 sequences from *Kluyveromyces lactis* (KLLA0_F22198g, KlBTA1), *Marchantia polymorpha* (MARPO_0023s0088, MpBTA1) and *Chlamydomonas reinhardtii* (CHLRE_07g324200v5, CrBTA1),were synthetized with codon optimization for expression in *A. thaliana* using the GeneArt synthesis service (ThermoFischer). The optimized sequences are provided in Supplementary Data 1. Forward/reverse primers used were atttcatttggagaggacacgctgacaagctgacATGTTCGGCTACATGAAGCACG/aagaagcgtagtcttgaacgtcgtatgggtaATTGTTCTCGAGGTCGAGTGAAG for KlBTA1, tcatttggagaggacacgctgacaagctgacATGGTGATGGAAAGCGTGAAGG/ aagaagcgtagtcttgaacgtcgtatgggtaCTTGGCATCGGTAGGAGTGC for MpBTA1 and ttcatttggagaggacacgctgacaagctgacATGGGATCTGGAAGAGATGGAAGG/ccaaagaagcgtagtcttgaacgtcgtatgggtaATTGTCCTTCTTAGCGCCCTTAC for CrBTA1. MgBTA1 gene (Naga_100016g36) was amplified from *Microchloropsis gaditana* cDNA with forward primer tttggagaggacacgctgacaagctgacatgtgcaatgttcctcgcgat and reverse primer agaagcgtagtcttgaacgtcgtatgggtactgggcttttgatttgccgg.

For PtBTA1-GFP construction, the coding sequence of *PtBTA1* gene (Phatr3_J42872) was amplified from *Phaeodactylum tricornutum* cDNA with forward primer gagaggacacgctgacaagctgactctagaATGATTGCCAGTCTCAAATCAC and reverse primer agtgaaaagttcttctcctttactggatccGTCCGTCTTTTTTTCTTCGCCC. For nf- PtBTA1-GFP construction, the *PtBTA1* gene (Phatr3_J42872) was amplified from *Phaeodactylum tricornutum* genomic DNA with forward primer catttggagaggacacgctgacaagctgacatgccggagatgcgttcccg and reverse primer aagcgtagtcttgaacgtcgtatgggtagtccgtctttttttcttcgccc. All amplified *BTA1* genes were then cloned in the pFP108 plant expression vector in frame with 3xHA tag by Gibson assembly (Gibson et al., 2009).

For MgBTA1-GFP and nf-BTA1Pt-GFP constructions, amplified *BTA1* genes were cloned in the pFP108 plant expression (To et al., 2006) vector in frame with 3xHA tags and GFP sequence in C-terminal and Sec63 (AtERDJ2B, AT4G21180) transmembrane fragment (aa 1-233) in N-terminal. For PtBTA1-GFP construction, amplified *BTA1* gene was cloned in the pFP108 plant expression vector in frame with GFP sequence in C-terminal (Figure S1). Expression of genes from this vector is under the control of two 35S promotors and a NOS terminator. Plasmids were transferred in *Agrobacterium tumefaciens* C58C1 strain and used for transient transformation of *N. benthamiana* leaves by agroinfiltration.

### Nicotiana benthamiana agroinfiltration

*A. tumefaciens* carrying BTA1 plasmids were cultivated in 10 mL of LB medium containing the appropriate antibiotics supplemented with acetosyringone (20 µM) during 24h at 28°C with 180 rpm agitation. *A. tumefaciens* strain expressing P19 protein was used in addition to all agroinfiltrations in the same conditions as *A. tumefaciens* transformants. Cultures were centrifuged for 10 min at 4000 rpm. The pellets were resuspended in 5 mL of agro-infiltration buffer (MgCl_2_ 1 mM, acetosyringone 200 µM) and further diluted to obtain 10 mL of cell suspensions at OD600nm=1. A mix of P19 and constructions volume to volume was used for infiltration with a 1 mL syringe at the abaxial side of leaves of 4 to 5 weeks old *N. benthamiana* plants. Plants were maintained for 4 or 8 days post-agroinfiltration and excised for microscopy and chloroplast purification or grounded in liquid nitrogen for western blot and lipid analysis.

### Confocal Microscopy

Laser scanning confocal microscopy was performed on a microscope Zeiss LSM880 equipped with a 63x/1.4 oil-immersed Plan-Apochromat objective, running Zen 2.3 SP1 acquisition software (Platform μLife, IRIG, LPCV). Excitation/emission wavelengths were 488/493 to 570 nm for GFP and 561/582 to 622 nm for RFP. Images were visualized and treated with the ImageJ 1.53a software running the image processing package Fiji. For the fluorescence quantification, 30 images per sample were analysed.

### Starch quantification

Starch quantification was performed with the Starch Assay kit from Sigma-Aldrich® (SA20). Leaves of *N. benthamiana* plants were harvested 4 and 8 days after agroinfiltration and directly frozen in liquid nitrogen. After grinding, the samples were supplemented with 5 mL of 80% ethanol and heated 5 min at 85°C in a water bath. The samples were mixed and supplemented with additional 5 mL of 80% ethanol before a 10 min centrifugation at 1000 x *g*. The following steps were performed following the manufacturer’s instructions. For the glucose assay, a standard curve was generated using 10, 20, 30, 40, and 50 µg of the kit’s glucose standard reagent and used to calculate the glucose quantity in µg/mg of fresh weight.

### Photosynthetic analysis

Chlorophyll fluorescence measurements were conducted on agroinfiltrated *N. benthamiana* plants at day 4 (four plants per measurements) at room temperature (RT) using a Speedzen 3 chlorophyll fluorescence imaging setup (JBeamBio, La Rochelle, France). Prior to analysis, samples were dark-adapted for 30 min. In this setup, actinic light and saturating pulses were generated by red LEDs peaking at 630 nm, while measuring pulses (duration of 250 ms) were emitted by blue LEDs peaking at 470 nm. The detection time following each measuring pulse was set at 15 μs, with no binning applied. The fluorescence value reported is an average obtained from 10 different points on various leaves, each consisting of four pixels. Maximal quantum efficiency of PSII was calculated as *F*_v_/*F*_m_f1=f1(*F*_m_ – *F*_o_)/*F*_m_, where *F*_o_ and *F*_m_ are the fluorescence emission measured before and at the end of a 200f1ms saturating pulse, respectively, in dark-adapted leaves. ΦPSII was calculated as (F_m’_f1-f1F)/F_m’_, with F being the steady-state fluorescence reached in presence of illumination with actinic light and *Fm’* being the fluorescence emission measured at the end of a 200f1ms saturating pulse, in the light-adapted leaves. The Electron Transport Rate (ETR) has been calculated as 0.6 x PAR x ΦPSII where 0.6 correspond to the average ratio of PSII reaction centers to PSI reaction centers and PAR is the irradiation light level in the PAR range (400nm to 700nm) in µmol photons.m^-2^.s^-1^. Nonphotochemical quenching (NPQ) was recorded during 5 min of illumination at 672 µmol photons.m^-2^.s^-1^ followed by 5 min of darkness. NPQ was calculated as (F_m_f1-f1F_m’_)/F_m’_(Maxwell and Johnson, 2000). Data were obtained from three independent experiments gathering three biological samples.

### Chloroplast purification

The agroinfiltrated plants were pre-incubated at 4°C for 12 hours prior to chloroplast purification. Leaves (3-4 g) were cut and homogenized in a blender in 70 mL ice-cold buffer containing 0.4 M sorbitol, 20 mM tricine/KOH pH8.4, 10 mM EDTA pH 8.0, 10 mM NaHCO_3_ and 0.15 % BSA (the supernatant was collected and stored at -80°C as the total fraction). A crude chloroplast pellet was obtained by centrifugation at 2070 x *g* for 2 min and further purified in 0.8 M sorbitol, 40 mM Hepes/KOH pH7.6, 10 mM MgCl_2_, 5 mM EDTA pH8.0, 20 mM NaHCO_3_ (washing buffer) on a Percoll gradient formerly prepared by centrigufation of 100% (v/v) Percoll in a JA20 rotor at 39191 x *g* for 55 min. After centrifugation at 9798 x *g* for 10 min, intact chloroplasts were retrieved from the lower phase. The chloroplasts were washed with 38 mL of washing buffer and centrifuged 5 min at 1482 x *g* in a JA20 rotor. The pellet was resuspended in 1 mL of washing buffer and stored at -80°C for further analysis as the purified chloroplasts fraction. The supernatant obtained after the first centrifugation was collected and centrifuged 1 hour at 4°C at 255530 x *g* in a SW40 rotor. The pellet was resuspended in 1 mL of washing buffer and stored at -80°C as the extraplastidial membranes fraction.

### Protein extraction and Western Blot Analysis

Proteins from *N. benthamiana* agroinfiltrated leaves were extracted from grounded tissue with 200 µL of protein buffer (urea 8 M, Tris-HCl 50 mM pH 6.8, EGTA 1 mM, DTT 10 mM) and proteins from chloroplast purification fractions were extracted with 500 µL of protein buffer. The proteins were heated 5 min at 70°C under 500 rpm agitation then quantified by Bradford assay (Invitrogen). 30 or 10 µg of proteins were used for migration on denaturing SDS-PAGE. After migration, the proteins were transferred on nitrocellulose membranes 1 hour at 25 Volt in transfer buffer MES SDS Running buffer 20x Bolt (Invitrogen). Membranes were stained with ponceau red (ponceau red 0.2% [w/v] and trichloroacetic acid 3% [v/v]) and used for western blots after destaining. Immunodetection was performed with anti-GFP-HRP (130-091-833, Miltenyi Biotec) and anti-BIP2 (AS09481, Agrisera) antibodies. Anti-LHCP antibody was kindly provided by Olivier Vallon (IBPC, Paris).

### Transmission Electronic Microscopy

*A. thaliana* leaves were prepared as previously described (Flori et al., 2018). Samples were fixed in 0.1 M phosphate buffer (PB) (pH 7.4) containing 2.5% (v/v) glutaraldehyde for 1 h at room temperature, under vacuum using a desiccator and then stored overnight at 4 °C. Samples were then washed five times in 0.1 M PB (pH 7.4), the buffer was removed directly by pipetting. Samples were fixed by a 2-h incubation on ice in 0.1 M PB (pH 7.4) containing 2% osmium and 1.5% ferricyanide potassium before they were washed five times with 0.1 M PB (pH 7.4). Samples were incubated in 0.1 M PB (pH 7.4) containing 0.1% tannic acid and incubated for 30 min in the dark at room temperature. Samples were again washed five times with 0.1 M PB (pH 7.4), dehydrated in ascending sequences of ethanol, and infiltrated with ethanol/ EMBed-812 resin mixture. Finally, the samples were embedded in EMBed-812. Ultrathin sections (70–100 nm) were prepared with a diamond knife on a PowerTome ultramicrotome (RMC products, Tucson, AZ, USA) and collected on 200 Mesh copper grids. Samples for Figure 7A and S8 were visualized by scanning transmission electron microscopy (STEM) using a FEI Tecnai microscope (FEI, USA) set up at 200 KV. For Supplementary movies and Figure 7B and C, focused ion beam (FIB) tomography was performed with a Zeiss CrossBeam 550 microscope (Zeiss, Germany), equipped with Fibics Atlas 3D software for tomography described previously (Uwizeye et al., 2021). The resin block containing the cells was fixed on a stub with silver paste, and surface-abraded with a diamond knife in a microtome to obtain a perfectly flat and clean surface. The entire sample was metallized with 4f1nm of platinum to avoid charging during the observations. Inside the FIB-SEM, a second platinum layer (1-2f1µm) was deposited locally on the analysed area to mitigate possible curtaining artefacts. The sample was then abraded slice by slice with the Ga^+^ ion beam (generally with a current of 700f1nA at 30f1kV). Each freshly exposed surface was imaged by scanning electron microscopy (SEM) at 1.5f1kV and with a current of ∼1f1nA using the in-column EsB backscatter detector. The simultaneous milling and imaging mode was used for better stability, with an hourly automatic correction of focus and astigmatism. For each slice, a thickness of 6f1nm was removed, and the SEM images were recorded with a pixel size of 6f1nm, providing an isotropic voxel size of 6 × 6 × 6 nm^3^. From the aligned FIB-SEM stack, the region of interest denoting the cell was cropped to remove unnecessary background using the Fiji software (https://imagej.net/Fiji). The 3D segmentation, reconstructions and visualization was achieved using Dragonfly software (www.theobjects.com/dragonfly). The surface areas and volumes were directly calculated from Dragonfly software.

### Lipid Analysis

Glycerolipids were extracted from plant samples frozen immediately in liquid nitrogen after harvesting. Once freeze-dried, the plant were crushed and suspended in 4 mL of boiling ethanol for 5 minutes to prevent lipid degradation and lipids were extracted according to (Simionato et al., 2013) by addition of 2 mL methanol and 8 mL chloroform at room temperature. The mixture was then saturated with argon and stirred for 1 hour at room temperature. After filtration through glass wool, cell remains were rinsed with 3 mL chloroform/methanol 2/1 (v/v) and 5 mL of NaCl 1% were then added to the filtrate to initiate biphase formation. The chloroform phase was dried under argon before solubilizing the lipid extract in pure chloroform. Total glycerolipids were quantified from their fatty acids: in an aliquot fraction, a known quantity of C15:0 was added and the fatty acids present were transformed as methyl esters (FAME) by a 1-hour incubation in 3 mL 2.5% H_2_SO_4_ in pure methanol at 100°C (Jouhet et al., 2003). The reaction was stopped by addition of 3 mL water and 3 mL hexane. The hexane phase was analyzed by gas chromatography coupled with a flame ionization detector (GC-FID) (Perkin Elmer) on a BPX70 (SGE) column. FAME were identified by comparison of their retention times with those of standards (Sigma) and quantified by the surface peak method using C15:0 for calibration.

The lipid extracts corresponding to 25 nmol of total fatty acids were dissolved in 100 µL of chloroform/methanol 2/1 (v/v) containing 125 pmol of each internal standard. Internal standards used were PE 18:0-18:0 and DAG 18:0-22:6 from Avanti Polar Lipid and SQDG 16:0-18:0 extracted from spinach thylakoid (Demé et al., 2014) and hydrogenated as described in (Buseman et al., 2006). Lipids were then separated by HPLC and quantified by MS/MS.

The HPLC separation method was adapted from (Rainteau et al., 2012). Lipid classes were separated using an Agilent 1260 Infinity II HPLC system using a 150 mm×3 mm (length × internal diameter) 5 µm diol column (Macherey-Nagel), at 40°C. The mobile phases consisted of hexane/isopropanol/water/ammonium acetate 1M, pH5.3 [625/350/24/1, (v/v/v/v)] (A) and isopropanol/water/ammonium acetate 1M, pH5.3 [850/149/1, (v/v/v)] (B). The injection volume was 20 µL. After 5 min, the percentage of B was increased linearly from 0% to 100% in 30 min and stayed at 100% for 15 min. This elution sequence was followed by a return to 100% A in 5 min and an equilibration for 20 min with 100% A before the next injection, leading to a total runtime of 70 min. The flow rate of the mobile phase was 200 µL/min. The distinct glycerophospholipid classes were eluted successively as a function of the polar head group.

Mass spectrometric analysis was done on a 6470 triple quadrupole mass spectrometer (Agilent) equipped with a Jet stream electrospray ion source under following settings: Drying gas heater: 230°C, Drying gas flow 10 L/min, Sheath gas heater: 200°C, Sheath gas flow: 10L/min, Nebulizer pressure: 25 psi, Capillary voltage: ± 4000 V, Nozzle voltage ± 2000. Nitrogen was used as collision gas. The quadrupoles Q1 and Q3 were operated at widest and unit resolution respectively. DGTS and PC analysis was carried out in positive ion mode by scanning for precursors of m/z 236 and 184 at a collision energy (CE) of 55 and 35 eV, respectively. SQDG analysis was carried out in negative ion mode by scanning for precursors of m/z -225 at a CE of -55eV. PE, PI, PG, PS, MGDG and DGDG measurements were performed in positive ion mode by scanning for neutral losses of 141 Da, 277 Da, 189 Da, 185 Da, 179 Da and 341 Da at CEs of 29 eV, 21 eV, 25 eV, 21eV, 8 eV and 11 eV, respectively. Quantification was done by multiple reaction monitoring (MRM) with 30 ms dwell time. DAG and TAG species were identified and quantified by MRM as singly charged ions [M+NH]^+^ at a CE of 19 and 26 eV respectively with 30 ms dwell time. DPG species were quantified by MRM as singly charged ions [M-H]- at a CE of -54 eV with 50 ms dwell time. Mass spectra were processed by MassHunter Workstation software (Agilent) for identification and quantification of lipids. Lipid amounts (pmol) were corrected by an in-house software developed by the LIPANG platform for response differences between internal standards and endogenous lipids and by comparison with a quantified control (QC). QC extract corresponds to a known mix lipid extract from Chlamydomonas and Arabidopsis cell culture qualified and quantified by TLC and GC-FID as described by (Jouhet et al., 2017).

### Statistical analysis

For photosynthesis measurements, normality was tested using Skewness and Kurtosis tests. Statistical differences between samples were analyzed using two-way ANOVA followed by Tukey’s multiple comparisons test.

For fluorescence quantification and lipid analysis, statistical differences between samples were analyzed using the pairwise comparison Kruskal-Wallis test followed by Benjamini, Krieger and Yekutieli post-hoc test.

## Acknowledgements

Sarah Salomon was supported by an INRAE PhD fellowship. Juliette Jouhet was supported by the Blink ANR (ANR-18-CE92-0015). The LIPANG (Lipid analysis in Grenoble) platform is supported by the Rhône-Alpes Region, the fonds FEDER, and GRAL, financed within the University Grenoble Alpes graduate school (Ecoles Universitaires de Recherche) CBH-EUR-GS (ANR-17-EURE-0003). We thank the microscopy facility MuLife of IRIG/DBSCI, funded by CEA Nanobio and GRAL LabEX (ANR-10-LABX-49-01) financed within the University Grenoble Alpes graduate school CBH-EUR-GS (ANR-17-EURE-0003). The authors would like to thank Sylvie Figuet for the maintenance of the plant growth facility and Dimitri Tolleter for his help for the photosynthesis measurement.

## Author Contribution

Sarah Salomon generated the cloning, the plant material and realized most of the experiments, Marion Schilling did the lipid analysis, Sylvaine Roy developed the softwares for reprocessing GC-FID and LC-MS data, Catherine Albrieux realized the chloroplast purification, Gregory Si Larbi, Denis Falconet and Pierre-Henri Jouneau achieved the electronic microscopy experiments, Morgane Michaud conceived the BTA1 expression in ER, the cloning strategy and stable transgenic *A. thaliana* lines, Juliette Jouhet designed and supervised the work and wrote the manuscript. All authors revised the manuscript.

## Supplementary Files

**Supplementary data 1**: Optimized sequences of BTA1 gene for plant expression.

**Supplementary movie 1:** FIB-SEM imaging from *N. benthamiana* leaves agroinfiltrated with MgBTA1-GFP. **Supplementary movie 2:** FIB-SEM imaging from *N. benthamiana* leaves agroinfiltrated with nf-PtBTA1-GFP. **Supplementary figures S1 to S9**

