## Supplementary data1 for "DGTS overproduced in seed plants is excluded from plastid membranes and promotes endomembrane expansion"

>KlBTA1-HA (codon optimized for expression in *A. thaliana*)

ATGTTCGGCTACATGAAGCACGTTGGGCAAGAGTGGAACGAGAAGATGCGTGAGTCTCCTAAGGTTGTGGGAGCTTCTGTGGGAATCATTACTCCTCTCGTTGTGCTCCTCATGTTCAGCGAGTCTCTCAGATCTGTTGTGCAGTTCTGTTGGCAGTGCTTCTTCAAGCCTTTCACCAGCGTGACCAGCAACAACAACCAGCAAGAAAACCTCGAGCAGTTCTACAAGAGCCAGGCTAAGCTCTACGATAGAACTAGGGGAGTTCTTCTGCAGGGGAGAGAGACTTCTCTTAAGCTCTCTCTCAGCCACCTCTCCGAGAAGAAGGGAAACGTTTGGATCGATGTTGGAGGTGGAACCGGGTTCAATATCAGCCAGATGGCTCTCCTTACCAACCTCGATACCACCTTCGACAAGATCTACCTGATCGACCTCTCTCCGTCTTTGTGCGAAGTGGCTAGAAAGAGGTGCAAAGAACACGGCTGGAAGAACGTTGAGGTGATCTGTGGTGATGCTTGCGACTTCGAGATCCCTGAAGAGTCTGCTCAGCTCATTACCTTCAGCTACAGCCTCTCTATGATCCCGTCTTTCTACGCTGCTATCGACCACGCTGTTTCACTCCTCGATGCTAAGAACGGCATCATCTCTTGCGTGGACTTCGGAGTGACTAACGAGTCTATGCTCGTTGGCAGAACTAACACTCTCGGAGGACTCGTGAACAGACACATCCCTTGGCTTTTCAGGACCTTCTGGCGTTTGTGGTTCGAGTTCGATAAGGTGTTCCTCGATCCTGCTAGACGTGAGTACCTCGAGTACAGATTCGGGACCATCAAGAGCCTCAACTGCTACAACTACAAGCTCGGGAAGATCCCGTACTACATCTGGCTTGGATGCAACAAGGATCACGAGCAGCATCTCCAGGCTCGTTTTGTTGAACTCGCTCCTACTTCTCCATACCTCGCTCCGATTACTACCAGCGCTTCTTCTAACGCTCAGCCGATGACTAAGGCTATGATCGCTGCTCTCGAGAACTCTAAGAAGGGACTCCCTTATCCGAGCCTCTTCTACCAGAAAGAGCATTGGAGGGTCTACTACGACGAGGTGAACCCTGAGTACAGCCAGTTCAAGAACAGCTACATCTACGCCTTCACCTGGGAAGATCCTCGTGAGGATGTGAACATCCTTAACATCCAGCCTGAGGACACGATCCTCGCTATTACTTCTGCTGGTGACAACATCCTCCACTACGCTACTCTTCCGAATCCTCCTAAGAGAATCCACGGCGTTGACTTGAACCCTTGCCAGGGACATCTTACCGAGTTGAAGCTTGCTGCTATCCGTTCGTTGTCTTTCACTCAGCTCTGGCAGATGTTCGGAGAGGGAAAGATCGACAGGTTCAACAACATCTTGCTCAACAAGCTCGCCCCGTACCTCTCTTCAAACGCTTTCCAATACTGGTTCGAGAACGGGACCAAGACCTTCGATCCTAATGGTGCTGGACTCTACGACACCGGATTCACTAAGTGGGCTCTTAGACTCGCTAAGTGGGTGTTCAAGGTGGCCAACCTCACTGATGAGGTTAACATGCTCTGCAAGGCTAAGACCCTTGAGGAACAGCGTTCTATCTGGGACAAGAAGATCAAGCCGGTCCTCTTCAACAGAGTGGTCGGAAAGATCCTTGTGGGGAACCCTCTTTTCCTTTGGTCTGCTCTTGGAGTGCCTAGAAACCAGGCAAAGATGATGGGATCTAGCACCCTCCAGTACATCATCGATACCCTCGATCCGGTGATCGACAACTCTCTCATCTCTGACGACAACTACTTCTACTACCTCACGCTCAAGGGCCGTTACTCTTCTAGATCTTGCCCGGACTACCTCAAAGAAGGCGGATTCAAGTCTCTGAGCAGGGAATCTCCTGAGTCTCCACTCGATAGAGTTAGGCTCCACACTGACACTCTCAAGGATGTTTGCGAGAGGCTCAGCAAAAAGACCGTGTCTATCGCCATCATCATGGACCACATGGATTGGTTCGATCCGCAGGGAACTGATGTGGATGAAGAGATCCAAGCTCTTTGGCTCGCTCTCAACTCTAGAGGAAGAGTTCTCCTCAGGTCCGCTTCTAAGTCTCCGTGGTACATCAAGAACTTCGAGAAGTTCGGGTTCAGCTGCAAGGCAGTGTCTGCTAGATATCCGGGAAAGTGCATCGACAGGGTGAACATGTATGCTTCTACTTGGGTGTGCCAGAAGATCAGCACTGCCTCTCAATCTAACAACCGTAGGCTCTCTTCACTCGACCTCGAGAACAATtacccatacgacgttcaagactacgcttctttgggtggttctagcccaagctcagagctccaccgcggtggcggccgcatcttttacccatacgatgttcctgactatgcgggctatccctatgacgtcccggactatgcaggagatctgtaa

>MpBTA1-HA (codon optimized for expression in *A. thaliana*)

ATGGTGATGGAAAGCGTGAAGGGGTACTACATGGATCTCCAGTGCCTCTCATCCATGTGGCTCAAGAAAATCAACGGCGGCTCTCACAAAGAGAGGCTCGAAGAATTCTACAGGCCTCAGGCTGAAGCTTACGATAGGTTCAGAGCTAACTTCCTCCATGGAAGGCAGCCTTTGCTTGCTGCTTGTGCTGCTAGACTCAGAGGATCTACTGGAATGATCTGGGTTGACCTCGGAGGTGGAACTGCTGAGAATGTGGATATGATGGCCCAGCTTATCGACCTCGCTAGCTTCGACAAGATCTACGTGGTGGATATGTGTGCGGCTCTCTGCAAAGTTGCTCGTGAGAAGGTTAGGCGTAAAGGCTGGTCTAACGTTGAGATCGTTGAGGGTGATGCCTGCACTTTCAAGCCTACTAAGCAGGCTACTCTCGTGACCTTCAGCTACAGCCTTTCTATGATCCCGACCTTCATGGATGCTGTGGACCAGTCTACTTCTTACCTCGCTGATGATGGAATGGTGGGGATCGCTGATTTCTTCACCTCTGCCAAGTACGATCTCCCTAACAGGCAGCATTCTTACGCCCAGAGATGGTTCTGGCGTACCGTGTTTGATTGCGACGGAATCGATGTTGGACCTGAGAGAAGGCAGTACCTCGAGCATAAGCTTGCTCCAGTGTTCGAGTTCAACGCTAGGGGTAAGATCCCTTACGTTCCGTACTTCCAGGCTCCGTACTACATCTGGATTGGAAGGCCTATCAAGTGCGACAAGGTGAACCTTCAGAGATTCCCTGCTAGGGCTTCTGATGAGTTTGAGGCTAAGAGGCCTCCTACTTTCCCTCCAACTTTCCTCTACTCGCTCTCTTGGGAAGATCCTCGTGAGGATGATAAGGTGCTCGAGATCAACCCTTCTGACACTGTTCTCACTCTCACCTCTGGTGGATGCAACGCTCTTGATCTCGTTTACCAAGGTGCTGGACAGGTTGTGGCTGTGGATATTAACCCTGCTCAGTCTTACCTCCTCGAGCTTAAGAGAGCTGCTGTTCTCAGACTCCCATTCGAGGATGTGTGGAAGATGTTCGGAGATGGTGTGCACCCTAAGTTCGAGCAGCTTCTCGATAGAGAGATCGCTCCATTCTTGTCTCAGGGCGCTGCTAACTTCTGGTATCAGAAGAGTTACTACTTCAAGAACGGGCTCTACTACCACGGTGGAATGGGTAGACTTCTCAGGGCTGTTAAGATCCTGGCTAAGGTGACCAGACAAGAGCATTGGGTTGACTGCCTTGTGAACGCTCCTACTCTCGAGAAGCAGAAAGAGCTTTGGTTCGCTACCGTTGGAAAGTGGCTTAGGCAGGCTAACATCTTCACCAGGATCAGCTCATTCCTCGTGACCAACAGACTCGTTCTTTGGTTCTGTGCTGGTGTGTGCAAGGGACAGTTGAGCCTTATCCGTAAAGAGGACAACATCTACGACTACATCGTGCGTTGCCTCAACTCTACCGCTGAGTACTCTCATCTCAGGGACAGCAACTACTTCTACAGATGCTGCCTCACCGGAAGGTTCTCTAGACAGTGTTGCCCTAGATTCCTTGAGCAGGGACCTTTCTACCAGCTCAAGGATGAACTTGCTAGGGGAGAGAGACTCCTCATCAGGACTGGAACTTTCGTGGGAGAGCTTAGGAAGAGGACCTACTCTAAGGTGATCCTCATGGATCACGTTGACTGGCTTGAGCAGCCTGATATCGATACCCTTTGTGACGCTCTCAGAGATCAGGTTAGGCCTGGTGGAAGAGTTATCTGGCGTTCTGCTTCAAGAAGGCCGGGATACGCTAAGTGCATTGAGAAGGCTGGATTCAAGGTGACCCGTATCCAGACCTCAGAGTCTTACATGGACCGTGTGAACATGTACGCCAGCTTCTATGTTGCCGTGAGGAACTCTGCTCCTTCTCCTATTCGTGTGCCTTCTGGAAACGTTGAGTCTACCGATGAGAGAAAGCCTCTCCTCTCTCCTGGATCTCCTGATTCTGTTCTCTGCACTCCTACCGATGCCAAGtacccatacgacgttcaagactacgcttctttgggtggttctagcccaagctcagagctccaccgcggtggcggccgcatcttttacccatacgatgttcctgactatgcgggctatccctatgacgtcccggactatgcaggagatctgtaa

>CrBTA1-HA (codon optimized for expression in *A. thaliana*)

ATGGGATCTGGAAGAGATGGAAGGCCTGCTAGCTACACCAAGAAGAACTTCTCACTCGAGAAGCTCAAGCTCAGCAGCATGAAGGATGATCTCACTGTGCTCAGGCACATGTGGTTCGGGTCTAAGAAAGGTGATGATCACGCTGCTAGGCTCGAGTCTTTTTACGGACCTCAAGCTGCTGCTTACGACGCTTTCAGATCTAGATTCCTCTGGGGACGTAGACCTATGCTTGCTGCTGTTGCTGCTAGACTTGCTGAGCGTTCTAACCTCATCTGGGTTGACCTTGGAGGTGGAACTGGTGAGAACGTTGACATGATGGCTGACTACATCGACCTCGCCAAGTTCAAGTCTATCTACGTGGTGGATCTCTGCCACTCTCTCTGCGAAGTTGCTAAGAAGAAGGCCAAGGCCAAAGGCTGGAAGAACGTTCAAGTTGTTGAGGCTGATGCTTGCCAGTTTGCTCCTCCTGAAGGTACTGCTACCCTGATTACCTTCAGCTACAGCCTCACTATGATCCCGCCTTTCCACAACGTTATCGACCAGGCTTGCTCTTACCTCTCTCAGGATGGACTTGTTGGAGTGGCCGATTTCTACGTGAGCGGAAAGTACGATCTTCCGCTCAGACAAATGCCTTGGAGCCGTAGATTTTTCTGGCGTAGCATCTTCGACATCGATAACATCGACATCGGACCTGAGAGAAGGGCTTACCTTGAGCAGAAGTTGGAGAGAGTTTGGGAGCAGAACACCCAGGGATCTATCCCTTATGTTCCTTGGCTTAGGGCTCCGTACTACGTTTGGATTGGTAGACTCCCTTCTGTGGGACATGCTCTCCATGAGGAAAGAGTGGAACGTCCTCCTATGTTCCCTCCTACTTTCCTGTACACCCAGAGCTGGGAAGATCCTGAGCCTGATATGGAAGTCATGGAAATCAACCCGAAGGACACCGTTCTCACTCTCACATCTGGTGGATGCAACGCTCTTAACCTCCTCGTTCAAGGTGCTGGACAGGTTGTGTCTGTTGATTGCAACCCTGCTCAGTCTGCTCTCCTCGAGCTTAAGAAAGTTGCTATCCAGCAGCTCGAGTTCGAGGATGTTTGGCAACTTTTCGGAGAGGGTGTGCACCCTAGAATCGAGGAACTTTACGAGAAGAAGCTCGCCCCATTCCTCTCTCAAACCTCGCATAACTTCTGGTCCAAGAGGCTCTGGTACTTCCAGCATGGACTCTACTACCAAGGTGGAATGGGAAAGCTTTGCTGGGTTCTCCAGTGCCTTGCTGTTGTTCTTGGACTCGGAAAGACCGTGAAGAGACTTGCTAACGCTCCGACCATGGAAGAACAGAGAAGGCTCTGGGATTCGAACATGCTCATCCACTTCGTGAAGAACGGACCTAAGCCTCTCGTTTGGCTGTTCGTGAAGTTCGTTTCTCTCGTCCTCTTCAACAAGGCCGTTCTTTGGTTTGGTGGTGGTGTTCCTGGAAAGCAGTACGCTCTTATCAAGGCTGACGGAATCCCGATCGAGAACTACATTGCTAGGACCATGGATGGTGTGGCCGAGAACTCTCATGTGAGGAAGCAGAACTACTTCTACTACAACTGCCTCACCGGGAAGTTCCTCAGAGATAACTGCCCTACTTACCTCCGTGAAGCTGCTTTCGCTACTCTCAAGTCTGGTGTGGTGGATAACCTCACCGTGAGCACCAATTTCTTCATGGAAGAGTTGAAGGCGAGGACCTACACGAAGGTGATCCTTATGGATCATGTGGACTGGCTCGATATGCCTGTGGCTAATGAGCTTGCTGAGTGCCTCGCTAAGCAAGTTGCTCCTGGTGGAATCGTTATCTGGCGTTCTGCTAGTCTCTCACCTCCTTACGCTGAGCTGATTCAAAAGGCTGGTTTCGACGTGAGATGCATCAGAAGGGCTACTCAGGGTTACATGGACCGTGTGAACATGTACTCGAGCTTCTACATGGCTAGGCGTAAGGGCGCTAAGAAGGACAATtacccatacgacgttcaagactacgcttctttgggtggttctagcccaagctcagagctccaccgcggtggcggccgcatcttttacccatacgatgttcctgactatgcgggctatccctatgacgtcccggactatgcaggagatctgtaa
