## Supplementary Figures S1 to S9 for "DGTS overproduced in seed plants is excluded from plastid membranes and promotes endomembrane expansion"

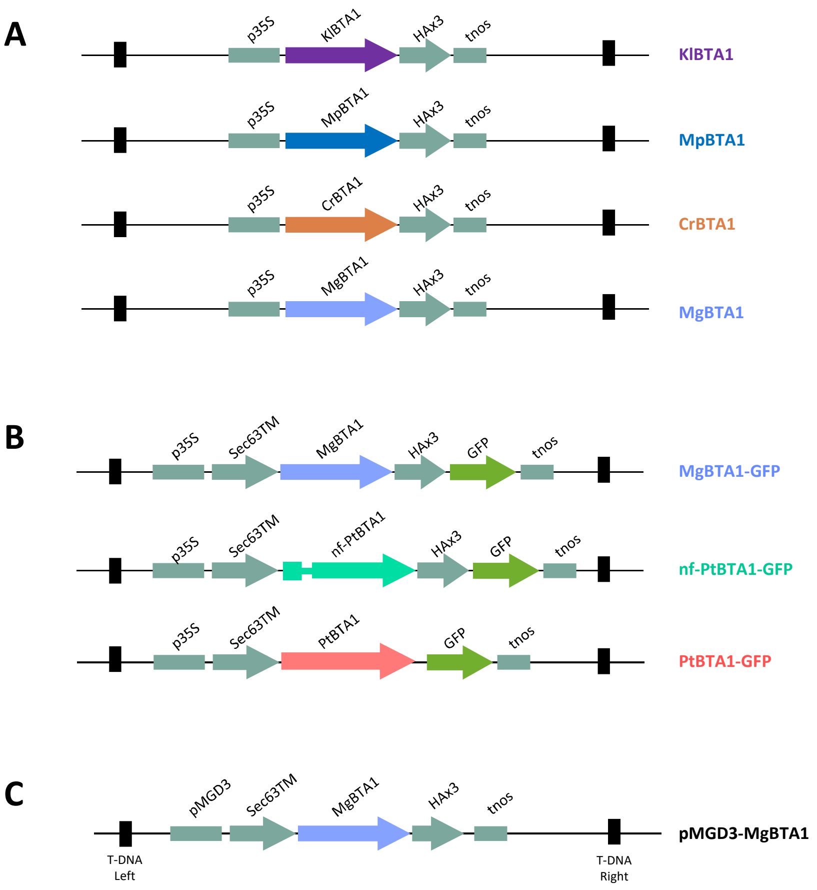


**Figure S1. Genetic constructions used for plant transformations**

(**A**) Schematic representation of the constructions used for *Nicotiana benthamiana* transient expression and subsequent lipid analysis for BTA1 activity shown in Fig 1. p35S : Cauliflower Mosaic Virus 35S promoter. KlBTA1, MpBTA1,CrBTA1 and MgBTA1 correspond to the *BTA1* gene from *Kluyveromyces lactis*, *Marchantia polymorpha, Chlamydomonas reinhardtii* and *Microchloropsis gaditana* (Naga_100016g36) , respectively

(**B**) Schematic representation of constructions used for *N. benthamiana* transient expression and subsequent analyses (lipid analyses, microscopy, chloroplast purification, western blot, starch quantification and photosynthesis analyses). MgBta1: BTA1 gene of *Microchloropsis gaditana* (Naga_100016g36); PtBTA1 : *BTA1* gene of *Phaeodactylum tricornutum* (Phatr3_J42872); nf-PtBTA1 : non-functional PtBTA1 protein containing an intron at the N-terminus of the protein; GFP : Green Fluorescent Protein.

(**C**) Schematic representation of the construction used for *A. thaliana* stable expression. MgBta1: BTA1 gene of *Microchloropsis gaditana* (Naga_100016g36); pMGD3 : MGD3 promoter inducible by phosphate starvation. Sec63TM : transmembrane fragment of Sec63 protein (targeting sequence to ER). tnos : nopaline synthase terminator.

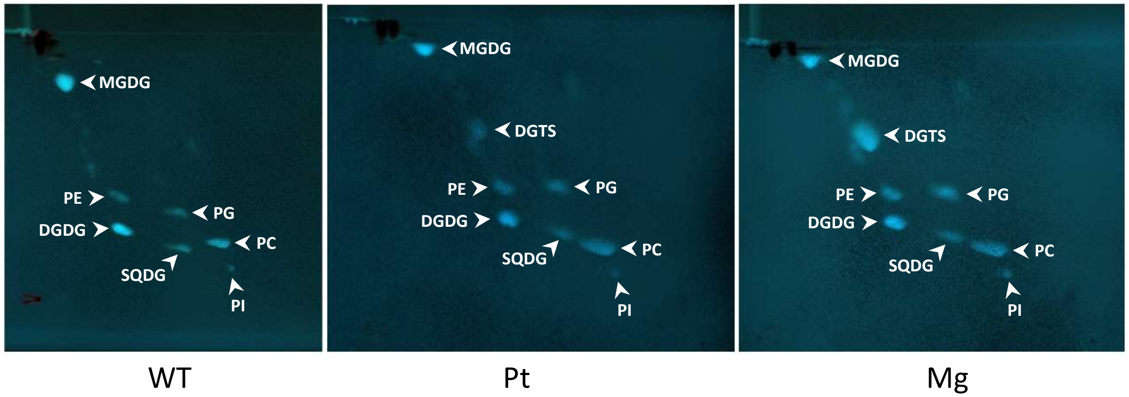


**Figure S2. 2D- Thin Layer Chromatography (TLC) analysis of lipid extracted from *Nicotiana benthamiana* leaves expressing BTA1 genes from *Phaeodactylum tricornutum* (Pt) and *Microchloropsis gaditana* (Mg).** WT : *Nicotiana benthamiana* leaves agroinfiltrated with P19 only. SQDG, sulfoquinovosyldiacylglycerol; MGDG, monogalactosyldiacylglycerol; DGDG, digalactosyldiacylglycerol; PG, phosphatidylglycerol; PI, phosphatidylinositol; PE, phosphatidylethanolamine; PC, phosphatidylcholine; DGTS, diacylglyceryl-N,N,N-trimethyl-homoserine.

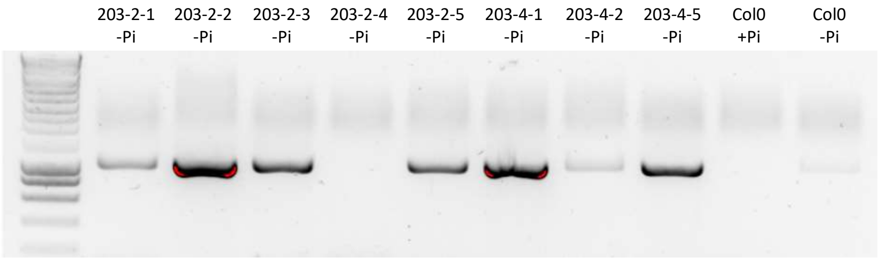


**Figure S3. Screening of *pMGD3-BTA1Mg-HA* seedlings**
PCR on Mg*BTA1* gene (Naga_100016g36) performed on gDNA of F2 seedlings lines grown in liquid MS medium depleted (-Pi) or repleted (-Pi) with Pi for Col0. Seedlings were harvested after 9 days of culture and grinded in liquid nitrogen. 2,5 µL of grinded seedlings were used for PCR analysis. 203-2-5 and 203-4-1 lines were kept and renamed pMGD3-MgBTA1-1 and pMGD3-MgBTA1-2, respectively.

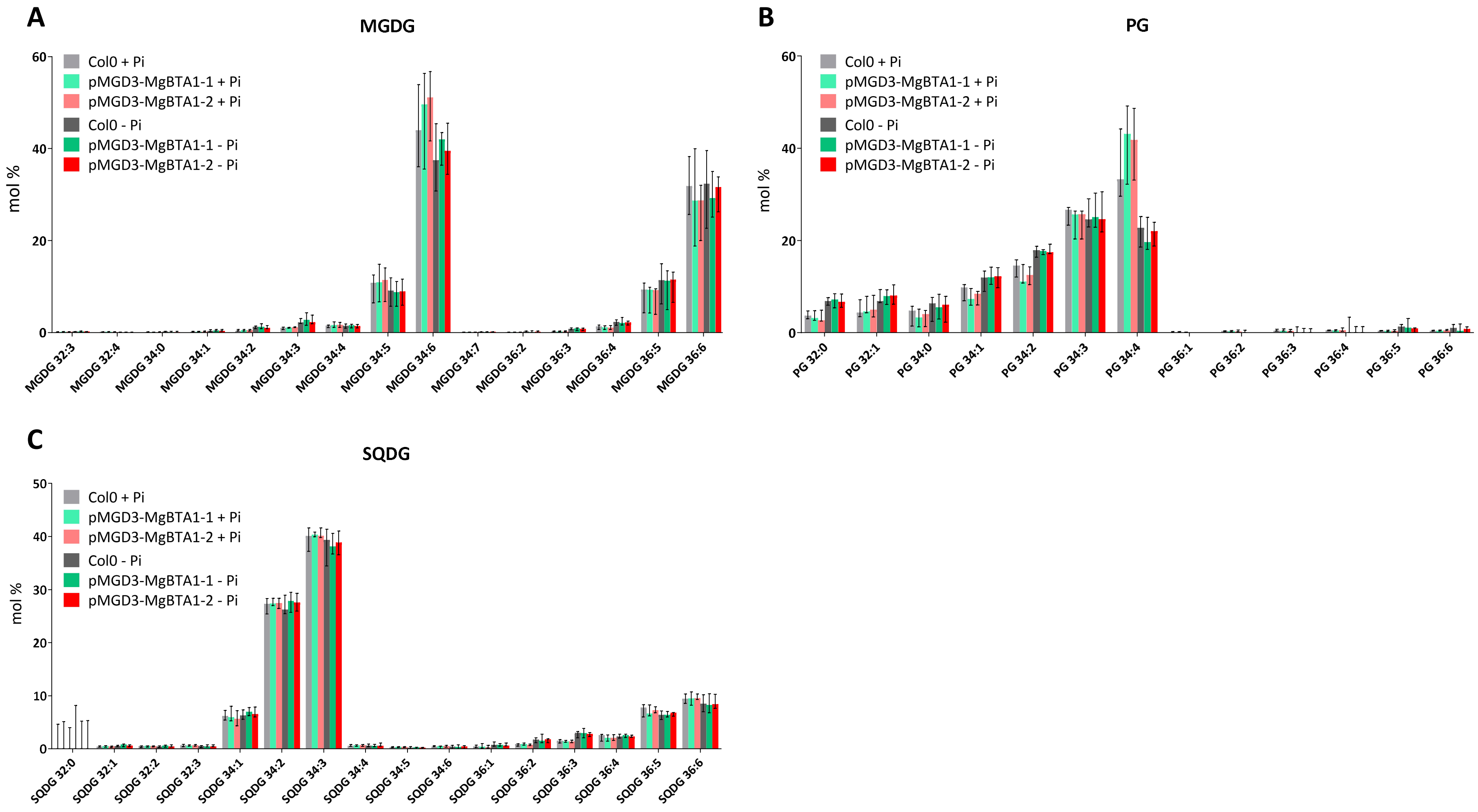


**Figure S4. Molecular distribution within chloroplast lipid MGDG (A), PG (B) and SQDG (C) classes determined by HPLC-MS/MS in *Arabidopsis thaliana* seedlings from Col0 and pMGD3-MgBTA1 lines n°1 and 2 grown 21 days in presence of 1 mM (+Pi) or 0,005 mM (-Pi) of phosphate. .** The first number corresponds to the sum of the carbon numbers of the two fatty acids, and the second number to the number of unsaturations. MGDG: monogalactosyldiacylglycerol; PG: phosphatidylglycerol; SQDG: sulfoquinovosyldiacylglycerol

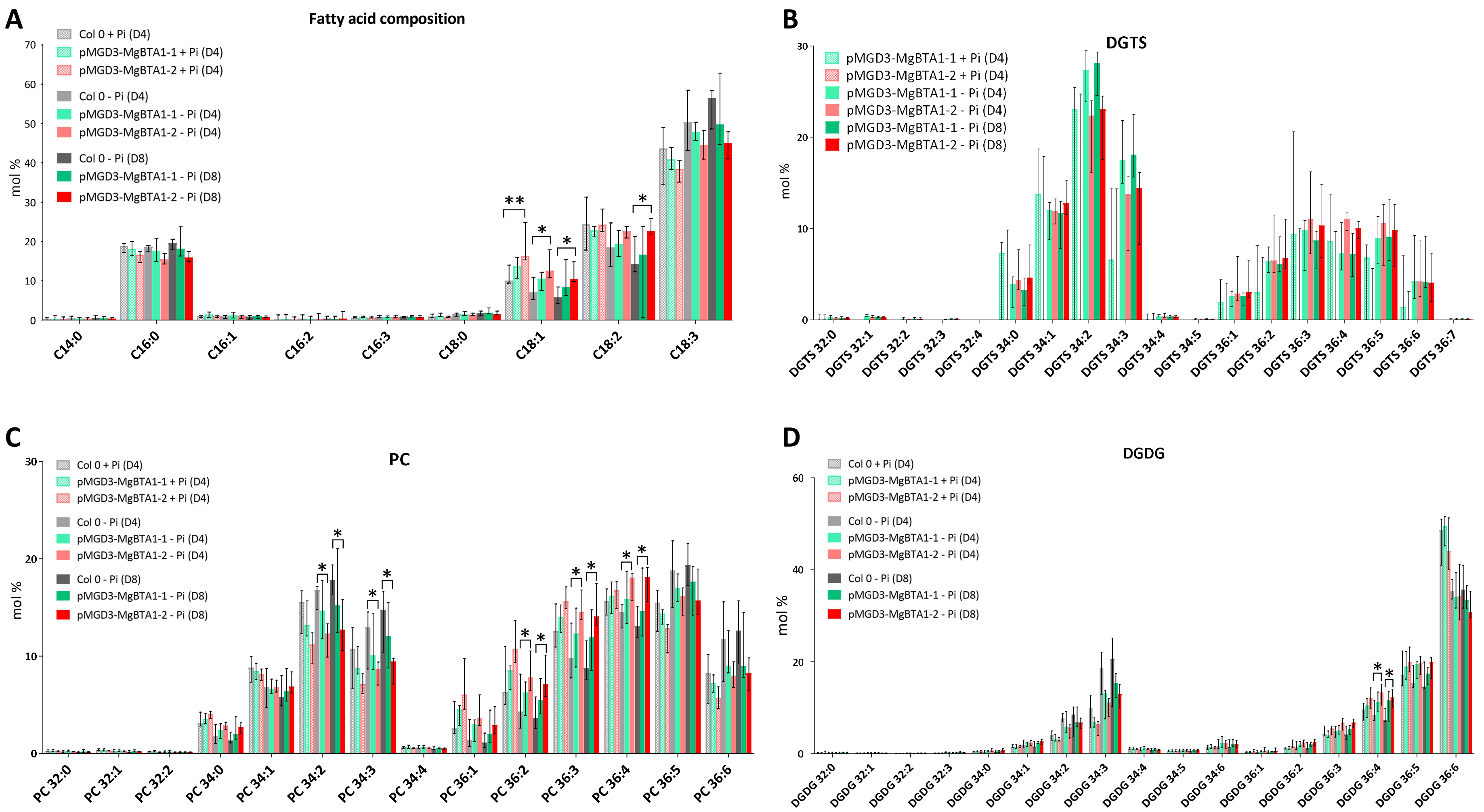


**Figure S5: Lipid analysis of *Arabidopsis thaliana* calli from Col0 and pMGD3-MgBTA1 lines n°1 and 2 cultivated 4 days in presence of 1 mM (+Pi (D4)) or 4 days (D4) and 8 days (D8) in absence of phosphate (-Pi) . (A)** Fatty acid distribution determined by GC-FID and **(B)**, **(C)**, **(D)** molecular distribution within DGTS, PC and DGDG classes respectively determined by HPLC-MS/MS. Asterisks indicate significant differences (***P*<0,01; **P*<0,05) in pairwise comparisons by a Kruskal-Wallis test.

DGDG, digalactosyldiacylglycerol; PC, phosphatidylcholine; DGTS, diacylglyceryl-N,N,N-trimethyl-homoserine. The first number corresponds to the sum of the carbon numbers of the two fatty acids, and the second number to the number of unsaturations.

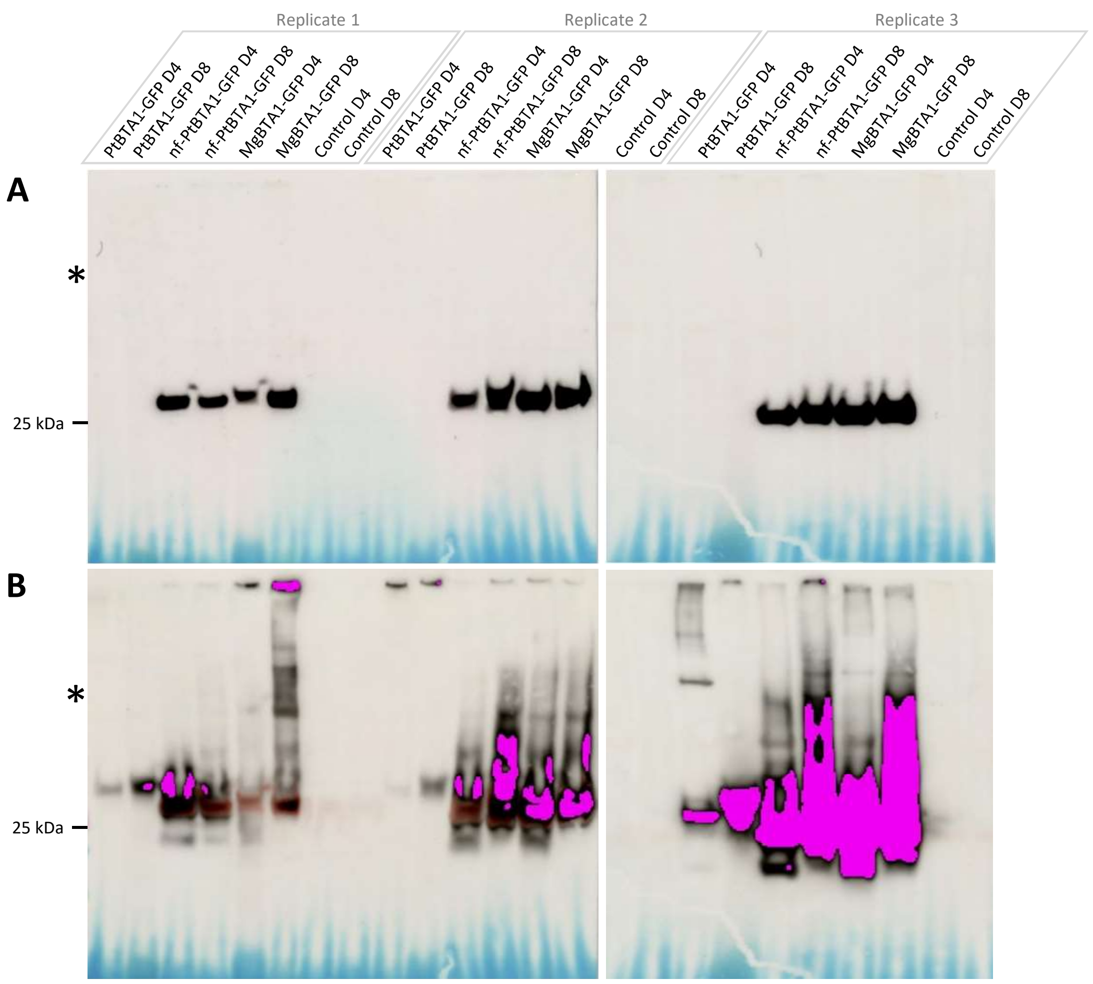


**Figure S6 Western Blot analysis of agroinfiltrated *N. benthamiana* leaves**

Western blot analysis of *N. benthamiana* leaves agroinfiltrated with BTA1 fused proteins (nf-BTA1Pt-GFP, BTA1Pt-GFP and BTA1Mg-GFP) and P19 using GFP antibody. Leaves were harvested 4 (D4) and 8 (D8) days after agroinfiltration. Asterisks indicate the expected size of the fused proteins (around 110 kDa). The GFP alone is at 27 kDa **(A)** Automatic exposure. **(B)** Exposure of 2 minutes and 20 seconds.

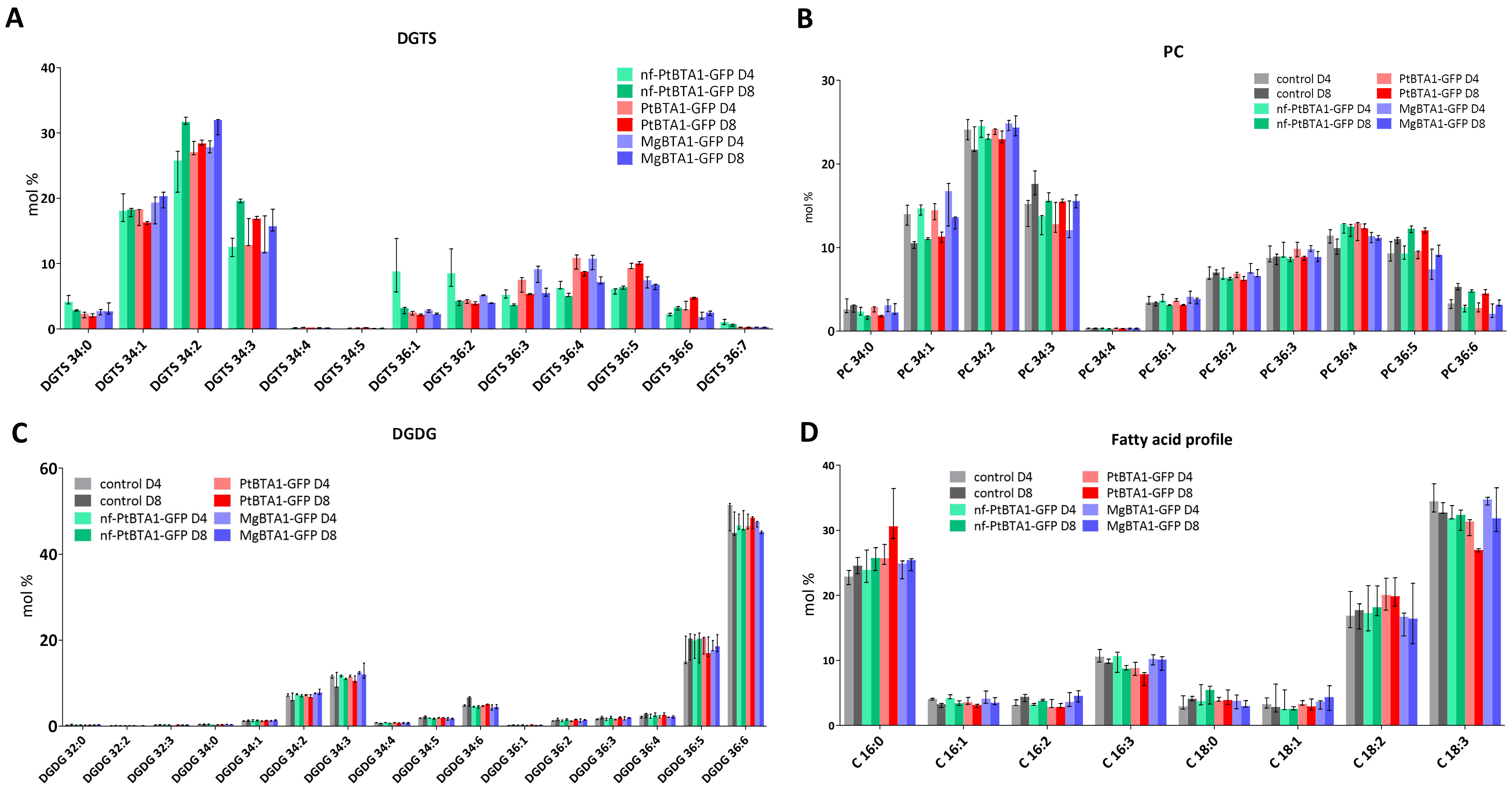


**Figure S7: Glycerolipid distribution of *Nicotiana benthamiana* leaves agroinfiltrated with or without (control) BTA1-GFP constructions.**

**(A)** to **(C)** Glycerolipid analysis and **(D)** fatty acid distribution of *N. benthamiana* leaves agroinfiltrated with P19 only (Control), nf-PtBTA1-GFP, PtBTA1-GFP and MgBTA1-GFP fused proteins. Leaves were harvested 4 (D4) and 8 (D8) days after agroinfiltration. Control: *N. benthamiana* agroinfiltrated with P19 protein (negative control). Represented values correspond to the median of 3 biological replicates and the error bars represent the range between minimum and maximum values. Molecular distribution within DGTS **(A)**, PC **(B)** and DGDG **(C)**. The first number corresponds to the sum of the carbon numbers of the two fatty acids, and the second number to the number of unsaturations. DGDG, digalactosyldiacylglycerol; PC, phosphatidylcholine; DGTS, diacylglyceryl-N,N,N-trimethyl-homoserine.

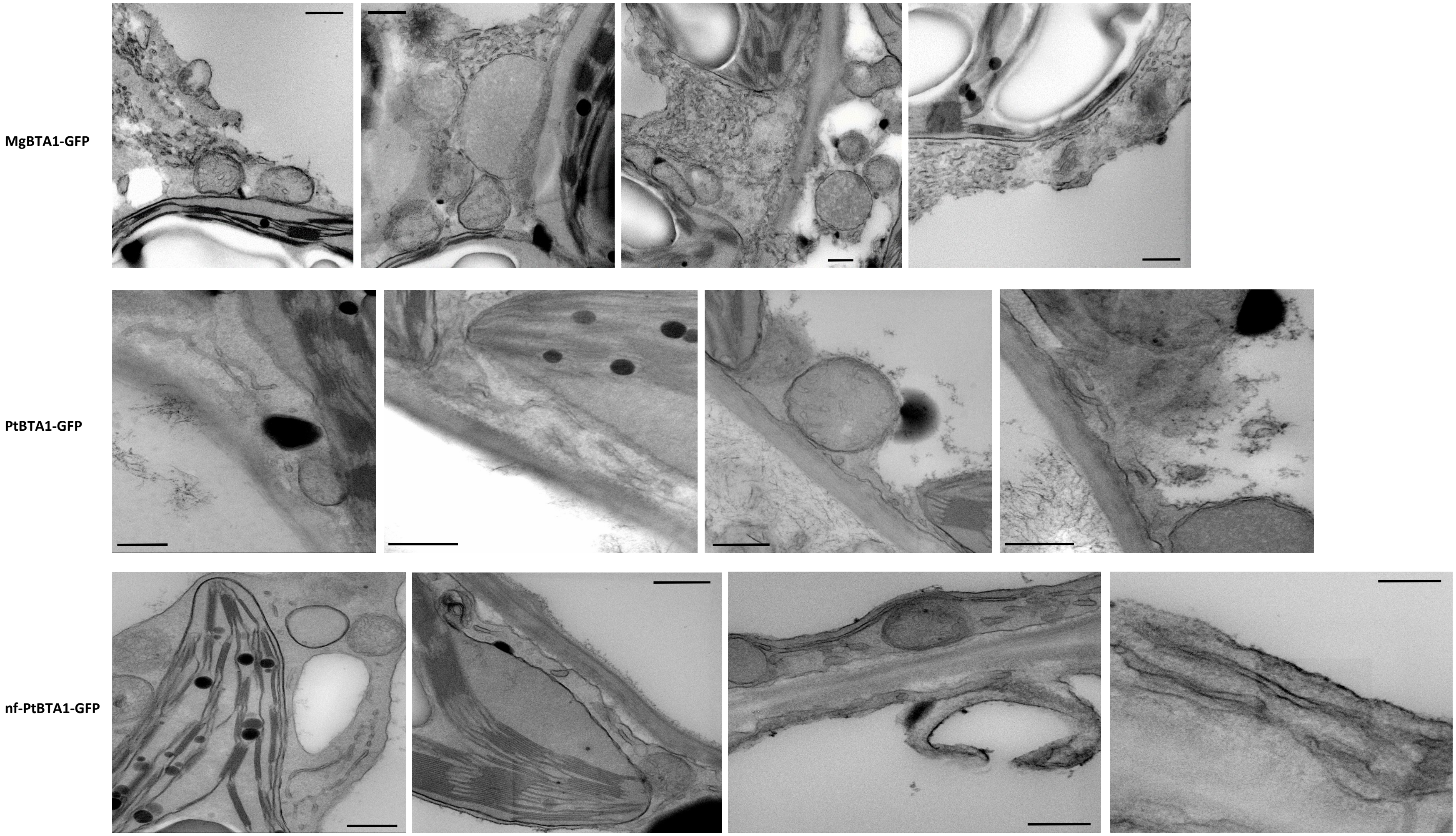


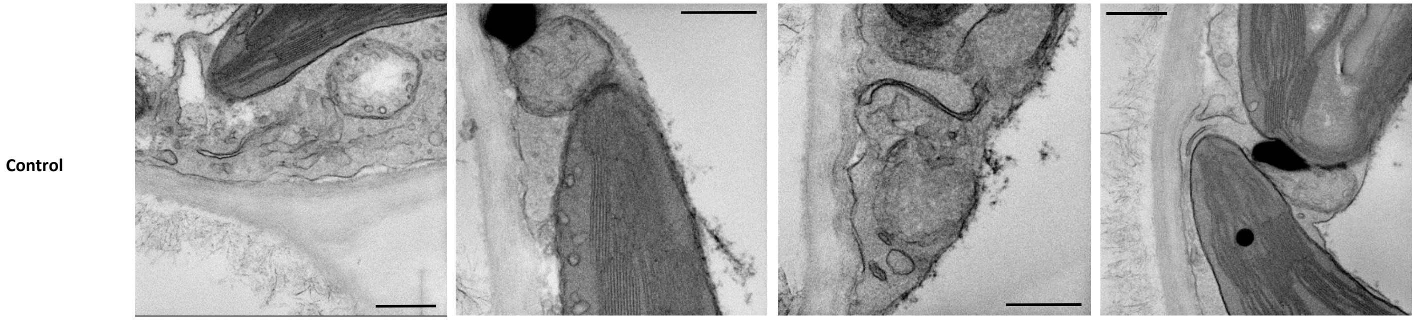


**Figure S8.** Electronic microscopy observation of the architecture of cellular membranes in *Nicotiana benthamiana* leaves agroinfiltrated with BTA1 fused proteins (nf-PtBTA1-GFP, PtBTA1-GFP and MgBTA1-GFP) or P19 as control at day 4. Scale bar : 500 nm.

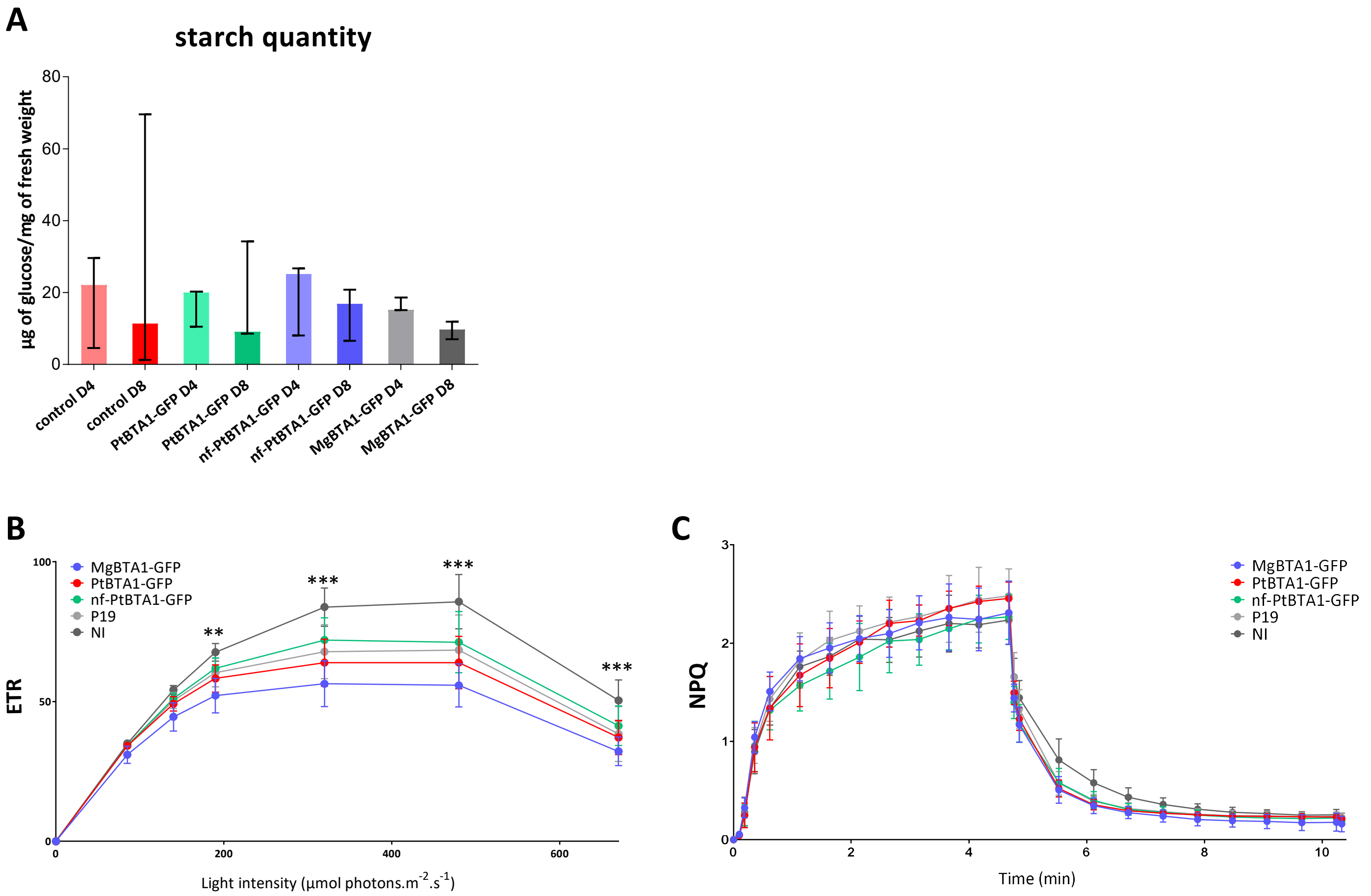


**Figure S9: Photosynthesis efficiency analysis on *N. benthamiana* leaves 4 days after agroinfiltration with BTA1-GFP constructions** (MgBTA1-GFP, PtBTA1-GFP, nf-PtBTA1-GFP, P19). NI : non infiltrated. P19 : *N. benthamiana* agroinfiltrated with P19 protein (negative control). Data are expressed as average values ± SD (bars) of three independent experiments.

**(A)** Starch quantification from *N. benthamiana* leaves agroinfiltrated with BTA1 constructions, 4 (D4) and 8 (D8) days after agroinfiltration.

**(B)** The Electron Transport Rate (ETR) has been calculated as 0.5 x PAR x ΦPSII where 0.5 correspond to the average ratio of PSII reaction centers to PSI reaction centers and PAR is the irradiation light level in the PAR range (400nm to 700nm) in µmol photons.m-2.s-1. Asteriks indicate significant differences (***P*<0,01;****P*<0,001) between MgBTA1-GFP and P19 samples by a Two-Way ANOVA followed by a Tukey's multiple comparisons test.

**(C)** Nonphotochemical quenching (NPQ) was calculated as (Fm - Fm')/Fm'. NPQ was recorded during 5 min of illumination at 672 µmol photons.m$^{-2}$.s$^{-1}$ followed by 5 min of darkness.
